# Targeted Genome Mining Discovery of the Ramoplanin Congener Chersinamycin from the Dynemicin-Producer *Micromonospora chersina* DSM 44154

**DOI:** 10.1101/2020.05.22.111625

**Authors:** Kelsey T. Morgan, Jeffrey Zheng, Dewey G. McCafferty

## Abstract

The availability of genome sequence data combined with bioinformatic genome mining has accelerated the identification of biosynthetic gene clusters (BGCs). Ramoplanins and enduracidins are lipodepsipeptides produced by *Actinoplanes ramoplaninifer* ATCC 33076 and *Streptomyces fungicidicus* B-5477, respectively, that exhibit excellent in vitro activity against a broad spectrum of Gram-positive pathogens. To explore if ramoplanin/enduracidin-like BGCs exist within genomes of organisms sequenced to date, we devised a targeted genome mining strategy that employed structure-activity relationships to identify conserved, essential biosynthesis genes from the ramoplanin and enduracidin BGCs. Five microorganisms were found to contain ramoplanin-like BGCs: the enediyne antibiotic producer *Micromonospora chersina* strain DSM 44151(dynemycin); the glycopeptide antibiotic producers *Amycolatopsis orientalis* strain B-37 (norvancomycin), *Amycolatopsis orientalis* strain DSM 40040 (vancomycin), and *Amycolatopsis balhimycina* FH1894 strain DSM 44591 (balhimycin); and *Streptomyces* sp. TLI_053. A single compound from fermentation of *M. chersina* was purified to homogeneity and found to possess good antibiotic activity against several Gram-positive bacterial test strains (1-2 μg/mL), comparing favorably to ramoplanin family members. We named this compound *chersinamycin* and elucidated its covalent structure, which differs distinctly from ramoplanins and enduracidins. Further, the chersinamycin BGC was validated through insertional gene inactivation using CRISPR-Cas9 gene editing. In addition to the information gained by comparing and contrasting the sequence and organization of these five new BGCs, the amenability of *M. chersina* to genetic manipulation provides a much-needed tool to investigate the fundamental aspects of lipodepsipeptide biosynthesis and to facilitate metabolic engineering efforts for the production of novel antibiotics capable of combating antibiotic-resistant infections.

## INTRODUCTION

For decades antimicrobial chemotherapy has been utilized successfully for the treatment of infectious disease. However, over the past thirty years, the rate of introduction of new-in-class antibiotics has flattened while the rate of clinical cases of infections due to bacteria that are resistant to front-line antibiotics has steadily increased, thus signaling a pressing need for the discovery and development of new antibiotic therapeutics.

Historically, natural products have helped meet this unmet need by providing a rich source of antimicrobial leads, as almost 70% of clinically approved antibiotics are natural products or second-generation natural product derivatives.^1^ For example, the glycopeptide antibiotics vancomycin and teicoplanin are first-generation natural products that have efficacy in their native form against infections from Gram-positive pathogens. Unfortunately, however, many first-generation natural products that possess good antimicrobial activity in vitro fail to make the jump to drug candidates because of several possible limitations: low stability, poor absorption, high toxicity, limited routes of delivery, and/or propensity to encounter resistance mechanisms. This creates a paradox in which these liabilities can preclude further investments in second-generation versions that could help overcome initial limitations, as exemplified by second-generation semisynthetic glycopeptides such as telavancin, oritavancin, and dalbavancin that exhibit markedly improved pharmacological properties and reduced toxicity profiles over the parent natural products.^2,3^

The ramoplanins are an exciting family of first-generation natural products that possess excellent in vitro activity against a wide range of Gram-positive bacteria. The family is composed of nonribosomally biosynthesized lipodepsipeptides that fall into two subclasses based on structure, the ramoplanins and the enduracidins (**Figure 1**). Ramoplanins, first isolated in 1984 by fermentation of *Actinoplanes* ATCC 33076,^4,5^ are a mixture of six lipoglycodepsipeptides of which factor A2 is most abundant, though all isomers possess similar antibiotic activities.^6,7^ The enduracidins A and B, lipodepsipeptides produced by *Streptomyces fungicidicus* B5477,^8–10^ are not glycosylated and contain longer N-terminal fatty acyl tails yet exhibit similar activity to ramoplanin. This antibiotic activity results from inhibition of bacterial cell wall biosynthesis. Ramoplanins and enduracidins capture peptidoglycan (PG) biosynthesis intermediate Lipid II, the substrate for transglycosylase and transpeptidase enzymes. Sequestering this late stage intermediate prevents formation of the mature, fully crosslinked peptidoglycan, resulting in a mechanically weakened cell wall and bacterial death due to osmotic lysis.^9,10^ In addition to interruption of PG biosynthesis, it has been reported that exposure of *S. aureus* to bactericidal concentrations of ramoplanin A2 results in membrane depolarization, suggesting a complementary mode of action through disruption of lipid membrane integrity.^11^

**Figure 1.**
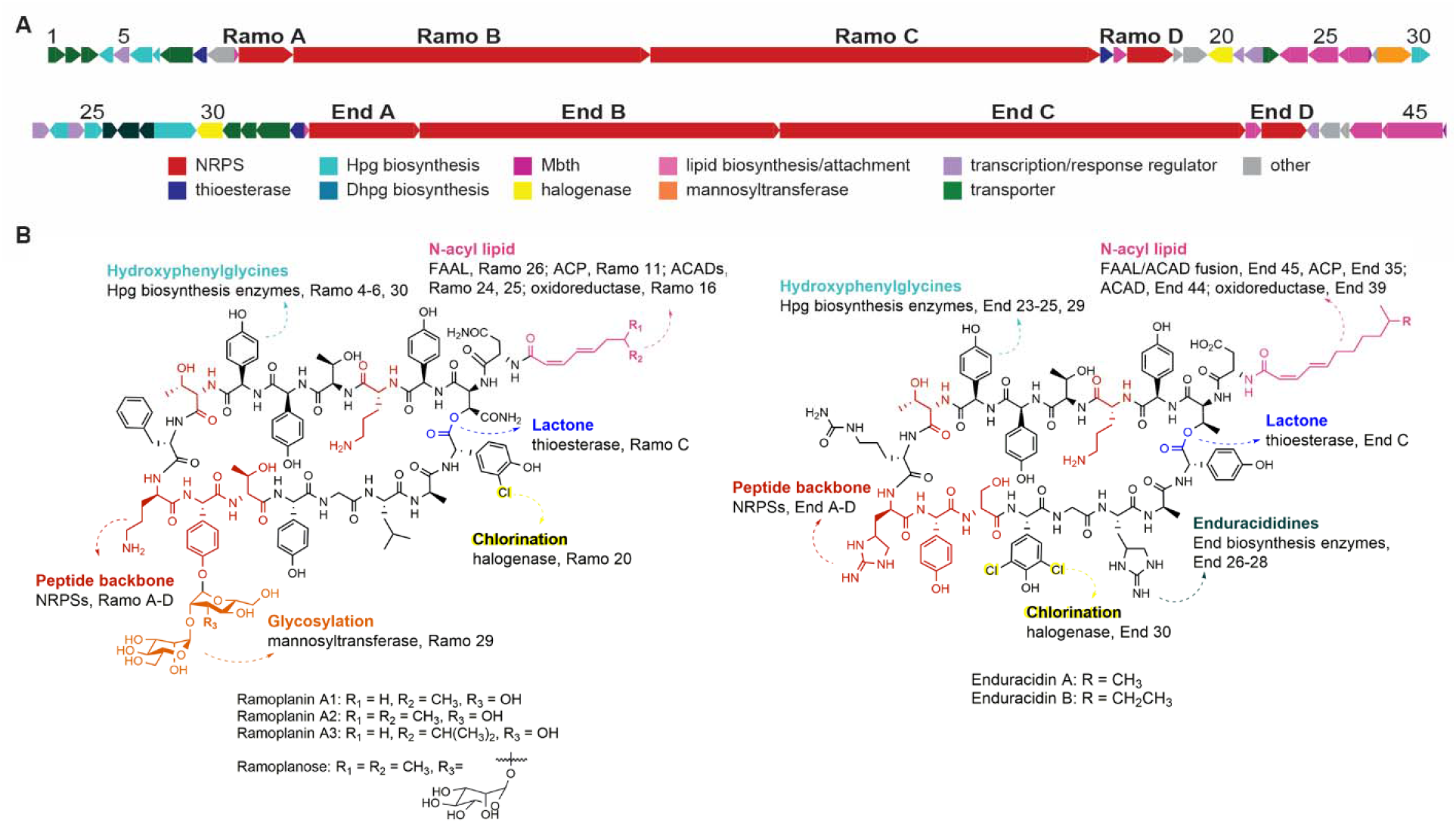
The ramoplanin family antibiotics. A) ORF arrow diagram depicting the defined BGCs from ramoplanin and enduracidin. B) Chemical structures of ramoplanin A2 and enduracidin A. Structural features are colored to coordinate with biosynthetic proteins responsible for their synthesis and incorporation, as shown in the ORF arrow diagram. “Hot spot” residues are in red and have been found to be essential by alanine scanning of ramoplanin, with MIC increases of >75-fold and K_d_ increases for substrate binding of >100-fold.

Ramoplanin A2 gained initial interest for treatment of Gram-positive bacterial infections that are resistant to antibiotics such as glycopeptides, macrolides, and penicillins.^9,12–15^ It has excellent in vitro activity with MICs ranging from 0.125–2 μg/mL. Unfortunately, this first-generation natural product could stand to be improved because it is not orally absorbed, is mild to moderately hemolytic when delivered intravenously, and its macrolactone is susceptible to hydrolysis when administered by intraperitoneal injection.^16^ Enduracidins A and B have a similar activity profiles, but exhibit reduced solubility and have been approved only for use outside of the United States as a growth-promoting feed additive for livestock.^17,18^

Despite minor limitations, ramoplanin was recently FDA approved for the treatment of *Clostridium difficile* colonic infections (CDI) and associated diarrhea.^19–22^ Oral delivery of ramoplanin achieves high colonic concentrations (>300 μg/mL), which far exceeds MICs determined in vitro against vancomycin-susceptible and vancomycin-resistant *C. difficile* strains (0.25–0.50 μg/mL). As such, the ramoplanin family of antibiotics remains a promising antibacterial agent warranting further development to broaden its therapeutic potential.

One underexplored avenue to develop second generation ramoplanin family members is to identify naturally produced congeners that may possess favorable structural diversities or allow for biosynthetic manipulations. In the case of glycopeptides, the development of second generation therapeutics has benefited from an arsenal of natural sources for investigation and development – there are over 30 glycopeptide-producing organisms giving rise to different core scaffolds and peripheral modifications such as acylation, glycosylation, and methylation that have provided insight into mode of action and been used to prioritize semisynthetic derivatization.^2,3^ Two recent studies have suggested the ramoplanin family of antibiotics are more widespread than previously assumed as well, and thus support such a strategy for accessing novel derivatives. First, during an unbiased screen of microbial extracts for activation of LiaRS reporter strains, de la Cruz and coworkers discovered 49 uncharacterized actinomycetes strains that produce compounds resembling ramoplanin A2 or ramoplanin analogs by LC/MS analyses.^23^ Independently, a second piece of evidence to this conclusion can be inferred from the recent work of Li and coworkers, who reported that enduracidin was produced by a second actinomycetes strain, *Streptomyces atrovirens* MGR140.^24^ These studies suggest that strains besides *Actinoplanes* and *S. fungicidicus* harbor biosynthetic machinery for ramoplanin congener production, and that new producing organisms could be uncovered to expand this important antibiotic class. As large scale sequencing efforts in recent years have generated a wealth of genomic data that can be mined with bioinformatics to identify biosynthetic gene clusters (BGCs)^25^ we envisioned that a genome mining approach could be used to identify new ramoplanin family producers.

In this study we developed a systematic method for uncovering ramoplanin-like BGCs within sequenced bacterial genomes. Previous structure activity relationship (SAR) studies of Reynolds, Boger, Walker, Ciabatti, and McCafferty groups were used to identify functionally important regions within the ramoplanin and enduracidin non-ribosomal peptide synthetases (NRPSs) and associated BGC standalone enzymes to serve as a suite of key sequence probes for genome mining.^15,16,26–35^ Using these SAR-informed protein sequences as search queries, we quickly developed a workflow that identified bacterial strains containing new lipodepsipeptide BGCs (**Figure 2A**). Herein we report the discovery of complete biosynthetic pathways for a ramoplanin family antibiotic in five new bacterial strains. Four of these five strains are host producers of either enediyne or glycopeptide antibiotics. One of these representative strains, the dynemicin producer *Micromonospora chersina* DSM 44154, was found to produce a ramoplanin congener we termed **chersinamycin**(**Figure 2B**). We report here the isolation, structure elucidation, antimicrobial activity, and validation of the BGC function using CRISPR-Cas9 gene editing of chersinamycin. These findings provide the foundation to further broaden our understanding of structure-function relationships among the ramoplanin family, to decode the molecular logic of ramoplanin biosynthesis, and to lay the groundwork for production of improved second generation ramoplanin analogs through mutasynthesis and metabolic engineering.

**Figure 2.**
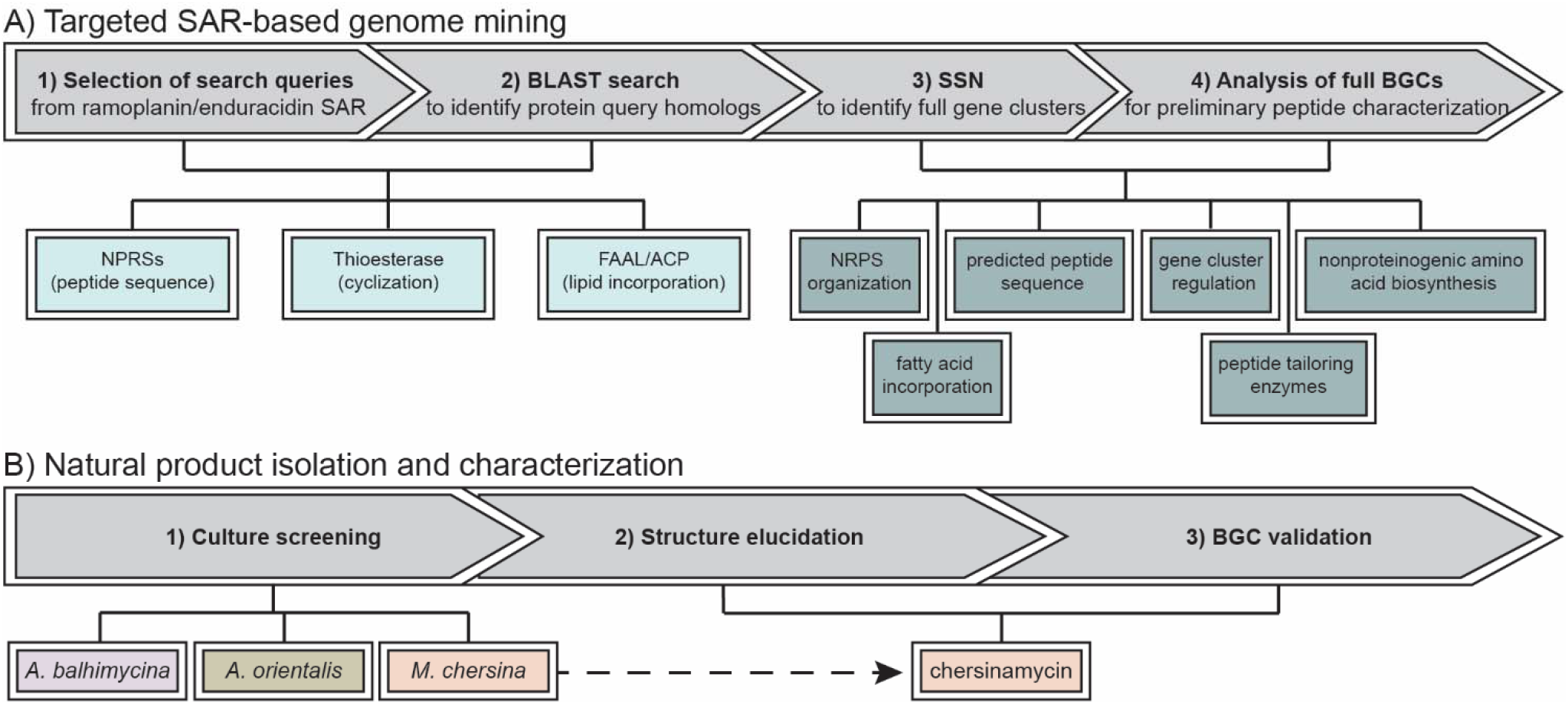
Workflow for the expansion of the ramoplanin family of antibiotics through targeted genome mining. A) Biosynthetic proteins and protein subdomains were selected from the ramoplanin and enduracidin BGCs and used as search queries for a targeted BLASTp search. Initial hits from the BLASTp search were moved forward to identify full gene clusters. B) Bacterial strains identified from SAR-based genome mining were screened for antibiotic production, with full characterization of the new lipoglycodepsipeptide chersinamycin from *M. chersina*.

## RESULTS AND DISCUSSION

### Examination of structure-activity analyses to inform genome mining rationale

The search for new ramoplanin family lipodepsipeptide gene clusters began with genome mining for key biosynthetic proteins linked to functionally important structural features, a process that was unique in that it was guided by results from structure-function studies of ramoplanins and enduracidins.^8–10,15,16,30–47^ There are several shared structural features of these antibiotics that are critically important for their activity: (1) Conserved amino acid type and stereochemistry within the 17-residue depsipeptide, which influences the overall peptide receptor-like conformation,^8,31,36,41^ promotes antibiotic dimerization,^31,37,47^ and facilitates binding to its lipid II target;^9,15,34,35^ (2) Conformational constraint imparted by the 49-atom macrocycle;^30,32–35,42,45^ and (3) N-terminal acylation, which promotes bacterial membrane association and influences its amphipathic C2 symmetrical dimeric conformation that is adopted upon membrane binding.^31,36^

Common to the ramoplanin and enduracidin BGCs are four NRPSs termed Ramo/End A–D (**Figure 1A**), which encode enzymes responsible for assembly line synthesis of these 17-residue peptides, including 12 nonstandard amino acids and seven with a D-amino acid configuration. Three large NRPS ORFs (*A*, *B*, *C*) appear to be organized in accordance with the collinearity rule of modular construction of NRPS condensation, adenylation, and thiolation domains. The exception is *ramoD*/*endD*, which encodes a standalone adenylation/thiolation di-domain enzyme that is predicted to work *in trans* with the NRPS B dual condensation/epimerization (C/E) domain to introduce D-*allo*-Thr^8^ within the linear peptide sequence.

Within the primary sequences of ramoplanin and enduracidin, there are several conserved residues that have been strongly linked to lipid II binding affinity and antibiotic activity. Boger and colleagues elegantly employed total solution-phase synthesis to perform an alanine scan of ramoplanin residues 3–13, 15, and 17 within [Dap^2^]-ramoplanin A2 aglycon, a hydrolytically stable ramoplanin aglycon analog.^30,32,34^ When compared to ramoplanin A1 A3 complex (MIC = 0.19 μg/mL), ramoplanin A2 aglycon (MIC = 0.11 μg/mL), and [Dap^2^]-ramoplanin aglycon (MIC = 0.07 μg/mL), alanine substitution of these 12 positions resulted in MIC increases over the parent antibiotics ranging from 1.3 to 540-fold (**Figure 1B**). Three positions exhibited markedly increased MICs: D-Hpg^3^ (74-fold), D-Hpg^7^ (53-fold) and D-Orn^10^ (540-fold). Residue 7 lies within the D-*allo-*Thr^5^-Hpg^6^-D-Hpg^7^-D-*allo-*Thr^8^ sequence that is conserved with enduracidins, and residue 10 is functionally conserved in enduracidins as D-enduracididine (End). Subsequently, Boger, Walker, and coworkers determined the effect of alanine substitution on lipid II binding and penicillin binding protein inhibition using a [Dap^2^]-ramoplanin A2 amide scaffold that was modified by the inclusion of single alanines along positions 3 12.^30,32^ The introduction of Ala residues increased *K_d_* values ranging from 378–8700 nM, with positions 4, 8, and 10–12 exhibiting >100-fold increased *K_d_*. Analogs that exhibited the most significant changes in MIC and *K_d_* values were considered functionally important and therefore likely to be conserved within a new ramoplanin/enduracidin congener. As such, these regions were carefully considered when devising our genome mining strategy.

The impact of conformational constraint imparted by the macrocycle was first investigated by Williams and coworkers, who demonstrated that hydrolysis of the macrolactone bond of ramoplanose resulted in a markedly less soluble linear peptide that lacked antimicrobial activity.^42^ In addition, Boger and coworkers showed that ramoplanin A2 activity required a 49-membered macrocycle, regardless of whether the macrocycle was linked by a lactone or lactam bond.^34^ Within Ramo C/End C NRPSs, the C-terminal thioesterase domain is responsible for installing this indispensable macrocycle^28,29^ and was considered a key biosynthetic sequence to be included as a genome mining search query.

Ramoplanins and enduracidins share genes that encode enzymes for fatty acid activation and lipoinitiation, the modification essential for bacterial membrane binding and antimicrobial activity.^28,29^ Both BGCs lack candidate ORFs encoding enzymes for *de novo* fatty acid biosynthesis, so it is likely that these fatty acids originate from primary metabolism and are activated as free fatty acids.^29,44^ In support of this hypothesis, an acyl carrier protein (ACP) and a fatty acid adenylate forming ligase (FAAL) appear in both BGCs. The presence of an N-terminal C^III^ condensation domain in NRPS A of both BGCs further supports a lipoinitiation mechanism involving fatty acid activation and condensation with residue one to form the starting N-acyl amino acid starter unit.

Although both antibiotic BGCs contain conserved acyl-CoA dehydrogenases (ACADs) and oxidoreductases that are believed to install the *E*,*Z* fatty acid double bonds, these enzymes are likely non-essential, since loss of these double bonds by hydrogenation of ramoplanin A2^48^ or semisynthesis^16^ resulted in no significant reduction in antimicrobial activity. Similarly, mannosylation^39,40^ and chlorination^43,49,50^ are structural elements that have been shown to be nonessential for antibiotic activity, although mannosylation has been shown by our group to enhance the conformational stability of ramoplanin A2^26,38^ and improve solubility over enduracidin.^37^

Collectively, these studies link membrane association, antimicrobial activity, and lipid II binding with specific structural elements shared between ramoplanin and enduracidin. By correlating functionally important architectural features with corresponding BGC-encoded enzymes that are responsible for their assembly, we identified a suite of probes for genome mining to search for new ramoplanin congeners.

### Discovery of ramoplanin-like biosynthetic gene clusters by genome mining

BGC sequences of seven SAR-guided probes from the NRPSs A–D, ACPs, and FAALs from the ramoplanin and enduracidin BGCs were used as initial BLASTp search queries to identify homologs from bacterial strains within the NCBI database. Protein sequence hits with >50% identity to the search queries were collected and cross-referenced to microbial strains that met the criteria of containing at least 4 homologs within its genome, regardless of ORF location. With these initial boundary conditions, 13 microbial strains were identified (**Table S1**). To determine if the protein homologs from the 13 strains were organized into a single BGC, we expanded the sequence analysis. Given the importance of the primary sequence encoded by the Ramo B/End B NRPS to the activity of ramoplanin and enduracidin, we analyzed the translated sequences within forty ORFs on either side of each NRPS B hit. Sequences obtained from the NCBI protein database were submitted to the EFI-Enzyme Similarity Tool^54^ for an all vs. all Blast search and assembly into a sequence similarity network (SSN) (**Figure S1**).

The SSN revealed clear protein clusters representing nearly all proteins within the defined ramoplanin and enduracidin BGCs; only five of the 24 proteins in the enduracidin BGC^32^ and six of the 31 proteins in the ramoplanin BGC^31^ are represented as isolated nodes. Though multiple proteins from each of the 13 preliminary strains were present within these clusters, we found that five strains contained all seven of the proteins utilized as genome mining probes localized to a single region of the genome. In addition, within the analyzed region of each of these five strains a significant number of ORFs were homologous to ramoplanin and enduracidin ORFs involved in nonproteinogenic amino acid synthesis, transcriptional regulation, and natural product transport. The strains found to encode a putative BGC for ramoplanin/enduracidin congener production include *Micronomonospora chersina* strain DSM 44151, *Amycolatopsis orientalis* strain B-37, *Amycolatopsis orientalis* strain DSM 40040, *Amycolaptopsis balhimycina* FH1894 strain DSM 44591, and *Streptomyces* sp. TLI_053 (**Figure 3**). Remarkably, four of these five new BGCs reside within bacterial strains that have been cultured and extracted for previously characterized natural products, including *A. orientalis* strain DSM 40040 and *A. balhimycina* FH1894, which produce the glycopeptide antibiotics vancomycin^52^ and balhimycin,^53^ respectively, and *M. chersina* DSM 44151, which produces the enediyne antibiotic dynemicin.^54^

**Figure 3.**
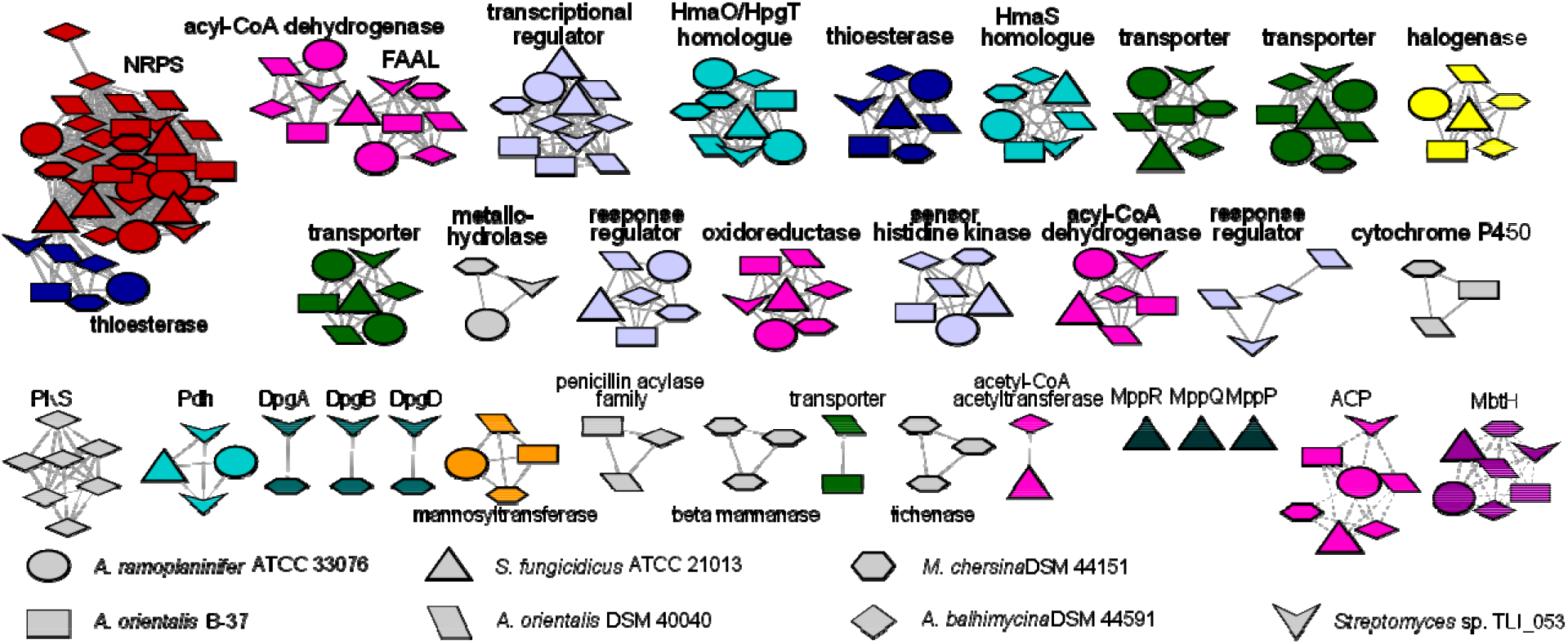
**Condensed sequence similarity network** for proteins within the BGCs of ramoplanin, enduracidin, and the five new ramoplanin family BGCs identified in this study. The network is assembled with an E value limit of 10^−5^ and alignment score of 50 (solid edges) or 25 (dashed edges).

The bounds of each of the five new BGCs were determined by analyzing clustered proteins within the SSN (**Figure 3, Figure S2A**). We identified remarkable similarity between ORFs included within the BGCs from each strain. The absence of clustered proteins not found within ramoplanin and enduracidin BGCs supports the previously defined bounds of the ramoplanin and enduracidin clusters.^28,29^ The gene organization and degree of conservation between each BGC likely reflects the necessity of nearly every protein in the cluster.

The SAR-guided genome mining approach allowed for the identification of five complete BGCs with strong similarity to the ramoplanin/enduracidin BGCs, suggesting that these five microorganisms contain the biosynthetic machinery to produce ramoplanin-like compounds. Manual analyses of increasingly stringent search criteria had the advantage of identifying candidates with inverted or varied organization of ORFs within the cluster, making them unable to be predicted by algorithms used by programs such as antiSMASH.^55^ This method was advantageous because it quickly allowed the selection criteria for hits to be filtered to select those most likely to belong to the desired antimicrobial class.

### In silico analysis of the NRPSs

Each of the five BGCs contained four NRPSs that are predicted to incorporate 17 amino acids into the peptide (**Figure S2B**). The organization of the NRPSs within each BGC was very similar to the ramoplanin and enduracidin NRPSs, including the presence of a standalone A-T domain of NRPS D, which suggests that these NRPSs also operate *in trans* with module 6 of each NRPS B that contains only C and T domains. NRPS A from each new cluster contains two full modules for the incorporation of two amino acids, leaving Ramo A as a unique NRPS in which a single module is predicted to act in an iterative fashion to assemble the first two asparagine residues.

The linear peptide sequence from each cluster was predicted from the adenylation domain specificity-conferring sequences.^56,57^ Web-based prediction software including NRPSPredictor2^58^ and the PKS/NRPS Analysis Web Site^59^ was complemented with manual sequence alignment of the eight conserved adenylation domain active site residues to account for genus-dependent sequence variation as well as a lack of predictive power for some unnatural amino acids by web-based software (**Table S3, Figure S2B**). For each organism, the NRPS-encoded primary sequences clearly predicted that all were likely ramoplanin congeners, yet each predicted sequence was unique and not identical to enduracidin or ramoplanin. Despite these differences, the NRPSs exhibited nearly identical conservation of five “hot spot” residues (Orn^4^, Thr^8^, Orn^10^, Hpg^11^, and Thr^12^) that had been identified as functionally important in ramoplanin for lipid II binding and antimicrobial activity. The only exception is residue 4 of the product encoded by the *Streptomyces* sp. TLI_053 NRPS, which predicts the ornithine is shifted to residue position 5 (**Figure S2B**).

Condensation domain sequences within the NRPSs were also examined using antiSMASH predictions and manual sequence alignment to identify C-domain subtypes (**Figure S3**).^60^ We found that each of the five organisms share a conserved starter condensation domain (C^III^) as the first domain of NRPS A for fatty acid incorporation at the N-terminal residue, consistent with the presence of a FAAL and ACP within the BGC and necessity of N-acylation for activity of ramoplanin and enduracidin. The order of classical ^L^C_L_ and dual C/E domains, responsible for incorporating L- and D-amino acids, respectively, exactly matches those found in the ramoplanin and enduracidin NRPSs within every module from the five new clusters (with D-amino acids in positions 3, 4, 5, 7, 8, 10, 12, and 16), with a single exception at NRPS A-module 2 of *M. chersina* and *Streptomyces* sp. TLI_053 NRPS A (**Figures S2B and S3**).

### Screening new bacterial strains for ramoplanin congener production

In an effort to identify and isolate new ramoplanin congeners, the three strains *M. chersina* strain DSM 44151, *A. orientalis* strain DSM 40040, and *A. balhimycina* FH 1894 strain DSM 44591 were examined for production of ramoplanin-like molecules. Initial media formulations screened included the optimized media for ramoplanin and enduracidin production,^44,61^ as well as the media optimized for production of each strain’s predominant natural product.^54,62–64^ Following incubation at various time intervals, cultures were extracted and screened by MALDI-TOF for a peptide within a mass range chosen based on bioinformatic predictions.

Although ramoplanin-like molecules were not observed to be produced by fermentation of either *A. orientalis* or *A. balhimycina*, fermentation of *M. chersina* for 12 days in dynemicin production medium H881 resulted in the production of a compound with a mass of 2574 Da that chromatographed similar to ramoplanin A2. This single compound was purified to homogeneity, generating yields of 1–3 mg/L (isolated, unoptimized yields). We named this compound **chersinamycin** and pursued its bioinformatics-guided structure elucidation and evaluation of its antimicrobial activity.

### In silico characterization of the chersinamycin BGC

To help reconcile the observed mass of chersinamycin with the predicted structure, we first examined the *M. chersina* strain DSM 44151 BGC, which is composed of 32 genes encoding proteins for transport, transcriptional regulation, amino acid biosynthesis, peptide assembly, and peptide tailoring (**Figure 4, Table S7**). In addition to the four NRPSs (Chers A–D) that are responsible for the production of a 17 residue linear peptide, the C-terminal thioesterase domain of Chers C suggests that the peptide is offloaded with concomitant cyclization (**Figure 4A, Figure S4**). While β-hydroxylation of the second amino acid, predicted as L-Asn, is difficult to predict based on adenylation domain sequence alone, a putative hydroxylase enzyme (Chers 38) was found in the chersinamycin BGC with high sequence identity to the ramoplanin hydroxylase (Ramo 10). A homologous enzyme is also identified in the *Streptomyces* sp. TLI_053 cluster, predicted to activate an aspartic acid at residue 2, but is absent in the additional four clusters which are each predicted to activate threonine at the second position (**Table S2**). Additionally, high percent identity between thioesterase sequences from the chersinamycin and ramoplanin clusters (**Figure S4**) suggested the site of macrolactonization to be the same.

**Figure 4.**
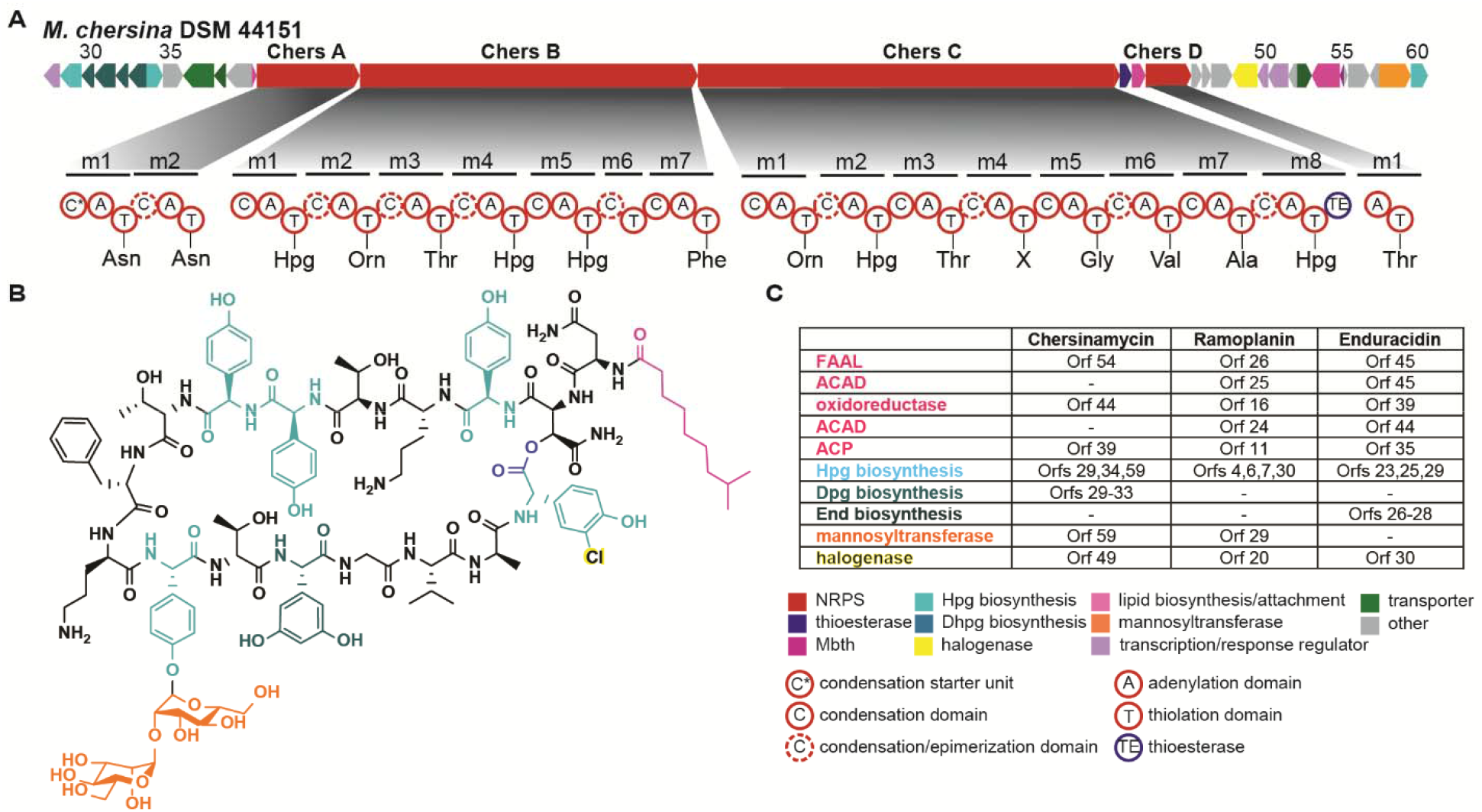
Structure and biosynthetic gene cluster of chersinamycin. A) ORF arrow diagram depicting the defined BGC from chersinamycin based on the generated SSN, and architecture of the four NRPSs within the chersinamycin BGC. Predicted amino acids based on adenylation domain specificity sequences are listed. B) Structure of chersinamycin as supported by bioinformatics and classical structure elucidation efforts. Structural motifs are colored according to the corresponding biosynthetic proteins responsible for their synthesis and incorporation. C) Comparison of biosynthetic enzymes found within the BGCs of chersinamycin, ramoplanin, and enduracidin.

Turning to the surrounding chersinamycin biosynthetic machinery, the presence of genes for Hpg biosynthesis (Chers 29, 34, and 59) supports the large number of predicted Hpg residues in the peptide sequence (**Figure 4A, Table S7**). At residues 4 and 10, the adenylation domain sequence confers specificity for a hydrophilic residue as predicted by NRPSPredictor2 (**Table S3**). The specificity sequences are nearly identical to those of ramoplanin and enduracidin at these positions, which contain Orn^4^, Orn^10^ and Orn^4^, End^10^, respectively. A lack of putative End biosynthesis proteins within the chersinamycin cluster led to the prediction of Orn^4^, Orn^10^ for chersinamycin.

Putative polyketide synthase-like (PKS-like) biosynthetic proteins Chers 29–33 with similarity to chalcone synthase and stilbene synthase suggested that chersinamycin may contain the amino acid 3,5-dihydroxyphenylglycine (Dpg).^65^ This amino acid is found within glycopeptides like vancomycin but absent in both ramoplanin and enduracidin. Though this residue was not directly predicted by NRPSPredictor2 or PKS/NRPS Analysis Web Site, an aromatic residue was predicted by NRPSPredictor2 at Chers C-m4 (residue 13). We predicted, therefore, that Dpg is incorporated at residue 13, and that Chers C may contain a novel Dpg-activating adenylation domain sequence.

As described previously, N-acylation has been determined essential to the antimicrobial activity of ramoplanin family antibiotics. In addition to the C^III^ domain of Chers A, a predicted FAAL (Chers 54) and ACP (Chers 39) are present within the cluster for fatty acid activation and transfer to the first NRPS-bound residue. Notably absent, however, was the prediction of putative ACADs (**Figure 4C, Table S2**). While an oxidoreductase is present (Chers 22), a lack of these dehydrogenases in the chersinamycin cluster suggests either a different biosynthetic source for an unsaturated lipid, or the incorporation of a saturated lipid.

Additional ORFs within the BGC appear to encode halogenase and glycosyltransferase tailoring enzymes. Chers 49 is homologous to the characterized halogenases found within the ramoplanin and enduracidin BGCs (Ramo 20 and End 30). Genetic knockout and complementation of Ramo 20 and End 30 within their respective clusters demonstrated that these enzymes are responsible for the monochlorination of Hpg^17^ in ramoplanin and dichlorination of Hpg^13^ in enduracidin.^43,50^ Identical adenylation domain specificity sequences at these sites and altered halogenation patterns resulting from genetic replacement of End 30 with Ramo 20 in *S. fungicidicus* suggested that site specificity of halogenation is controlled by the local structural environment of the full peptide, rather than loading of a halogenated residue onto the NRPS. We were therefore unable to confidently predict the location of possible halogenated residues for chersinamycin, but the high sequence similarity of Chers 49 to Ramo 20 and End 30 led us to suspect chlorination of an aromatic residue. Finally, the chersinamycin BGC contains a putative mannosyltransferase, Chers 59. The ramoplanin mannosyltransferase Ramo 29 has been implicated through genetic knockout and complementation to instill two D-mannose sugars onto the phenolic oxygen of Hpg^11^,^39,49^ and therefore glycosylation was predicted for chersinamycin as well.

### Chersinamycin isolation and structure elucidation

Numerous analytical methods were employed for the full structure elucidation of chersinamycin. HR-LC/MS revealed a [M+2H]^2+^ molecular ion of 1287.0511, suggesting a molecular formula of C_119_H_158_ClN_21_O_41_. The peptide macrocycle was determined to be highly base labile, with exposure to 1% triethylamine in water resulting in hydrolysis ([M+2H]^2+^ molecular ion 1296.044). This suggested a lactone macrocycle as opposed to a lactam which would remain intact under such weakly basic conditions, supporting our prediction that ring closure occurs at a side chain hydroxyl.

The ^1^H-NMR of the cyclic peptide was first examined to assign individual residues. A large number of exchangeable amide protons (δH 7.0–10.0) and signals within the α-proton region (δH 3.5–7.0) were observed, as well as many doublets in the aromatic region consistent with numerous Hpg residues (δH 6.0–7.5). Analysis of 2D NMR data allowed the assignment of the 17 amino acid residues (**Table S4**). COSY and TOCSY correlations were used to assign full aliphatic residues, confirming the incorporation of valine, alanine, glycine, threonines and ornithines into the peptide. COSY correlations between aromatic resonances in conjunction with NOEs between these resonances and their amide and alpha protons allowed the assignment of full aromatic residues. Two diagnostic singlets at δH 6.04 and δH 6.09 suggested a Dpg residue, supporting our predictions based on the Dpg biosynthetic proteins within the gene cluster. Correlations observed between several resonances in the region between δH 3.0–5.0 are consistent with the presence of sugar moieties which we hypothesize to be incorporated by Chers 59. Though exact resonances could not be assigned due to spectral overlap, resonances were identical to those observed in ramoplanin, which coupled with the presence of a putative mannosyltransferase within the BGC suggest D-mannoses are incorporated.

The N-acyl lipid was first assigned by 1D and 2D NMR. The 1D ^1^H NMR displays a strong doublet at δH 0.65 indicating a terminally branched lipid. Unlike the diagnostic spectra for the *Z,E* unsaturated lipids of ramoplanin and enduracidin, the ^1^H-NMR of chersinamycin showed a lack of vinylic protons, and 2D spectra lacked correlations spanning the aliphatic-to-olefinic region, supporting our hypothesis of a saturated lipid based on the lack of ACADs in the gene cluster. To confirm saturation, chersinamycin was additionally subjected to catalytic hydrogenation. While hydrogenation of ramoplanin reduces both olefins resulting in a mass increase of 4 Da, no change was observed for chersinamycin after 24 hours under hydrogenation conditions.

After defining the amino acid, sugar, and lipid components of chersinamycin, the peptide connectivity hypothesized from in silico analysis of the chersinamycin NRPS domains was further examined through analysis of the NOESY spectrum. NOEs between adjacent amide protons and between amide protons and adjacent alpha/beta protons allowed for connectivity to be determined (**Figure S19**). Strong NOE correlations between residues 2 and 17 supported macrolactonization between these residues as had been predicted through bioinformatics. To further validate connectivity, we turned to MS/MS. Fragmentation focused on the molecular ion [M+2H]^2+^ (1287.05) resulted in two highly abundant doubly charged product ions of 1206.013 and 1124.986, each consistent with a loss of a mannose residue from the core peptide. Unfortunately, the high fragmentation energy required to fragment the peptide resulted in many ions that were not diagnostic, a common occurrence with cyclic and glycosylated peptides. MS/MS of acyclic chersinamycin focused on the molecular ion [M+2H]^2+^ (1296.04) resulted in a more simplified spectrum (**Figures S11, S12**). Assignment of a number of b- and y-ions validated that hydrolysis occurred between residues 2 and 17, and confirmed the connectivity shown in **Figure 5**.

**Figure 5.**
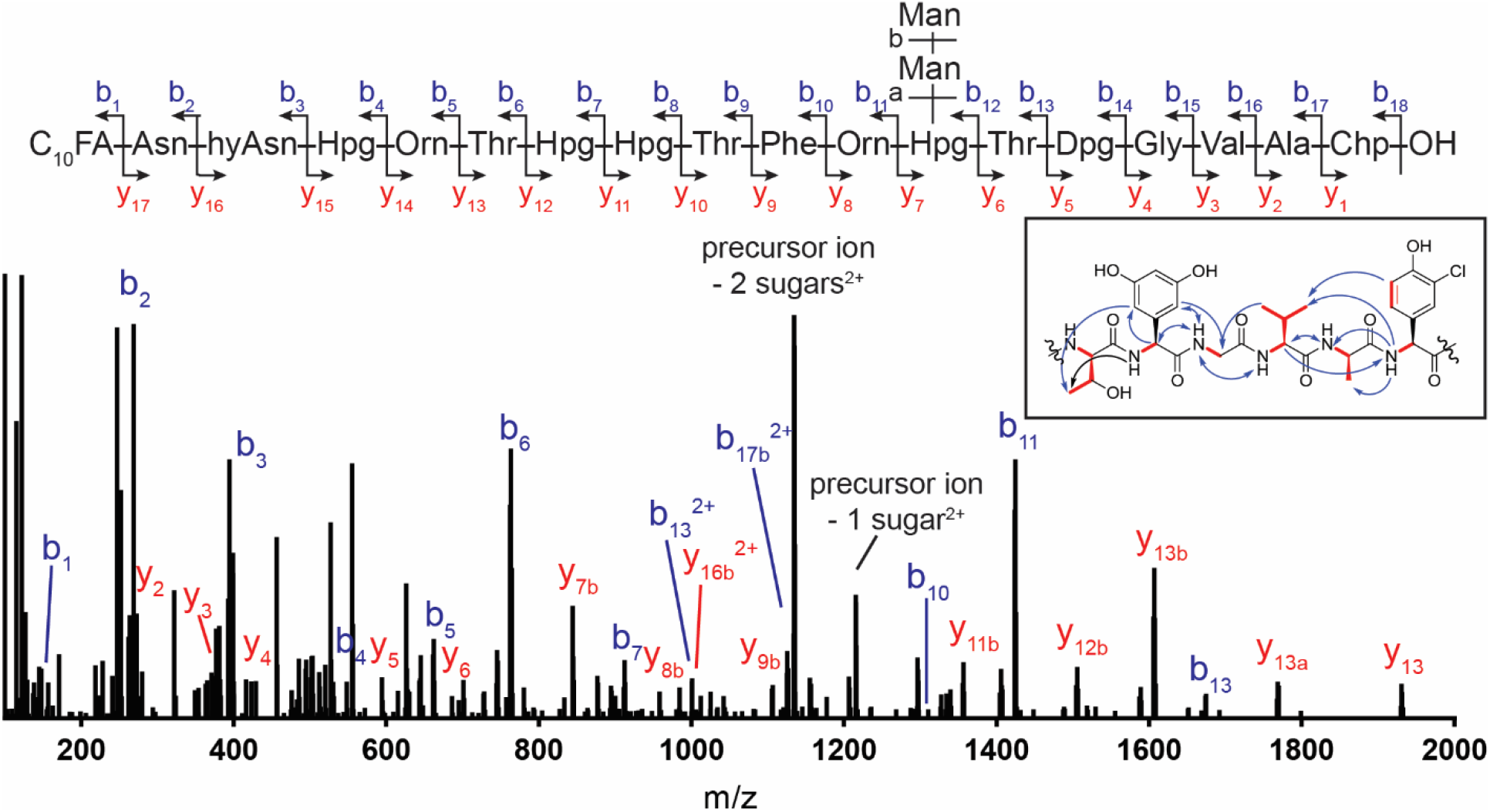
Structural characterization of chersinamycin. The MS/MS spectrum of acyclic chersinamycin is shown with the diagnostic fragmentation pattern of b- (blue) and y-ions (red). Inlaid figure shows COSY/TOCSY (red) and NOESY correlations (blue) for a key region of Dpg13-Chp17, which differs significantly from ramoplanin. Tables of calculated and observed MS/MS ions and full COSY and TOCSY spectra are included in the Supplemental Information.

Advanced Marfey’s analysis was employed to confirm the absolute configuration of each amino acid. Following complete hydrolysis and derivatization with Marfey’s reagent (FDAA), the hydrolysate of chersinamycin was analyzed by LC-MS and peaks were compared to authentic standards of FDAA-amino acids (**Figure S13**). It was determined that alanine and both ornithines are D-amino acids and valine, phenylalanine, and chlorohydroxyphenylglycine are L-amino acids. We observed a 1:1 ratio of D-Hpg:L-Hpg. Our chromatography method was able to unambiguously distinguish D/L-Thr from D/L-*allo*-Thr, allowing us to assign all threonines in chersinamycin as D-*allo*- and L-*allo*-Thr. The positions of D/L-amino acids in which both stereoisomers are present were assigned based on our analysis of the NRPS C/E domains. Unfortunately, asparagine and dihydroxyphenylglycine could not be identified in our FDAA-hydrolysate. As such, we were unable to confirm absolute configuration of these residues and assigned stereochemistry is based on the presence or absence of C/E domains.

Cumulatively, our bioinformatics analyses paired with analytical structure elucidation assigns our 2574 Da peptide from *M. chersina* as a 17-amino acid cyclic lipoglycodepsipeptide. The presence and location of D- and L-amino acids suggests chersinamycin’s 3D structure to be very similar to ramoplanin and enduracidin. Unique from ramoplanin and enduracidin, chersinamycin exhibits a saturated N-acyl lipid and a noncanonical Dpg residue within the peptide sequence. The observation of glycosylation is an advantageous structural feature for solubility, stability, and possible drug development. With the structure elucidated we next aimed to unambiguously confirm the BGC and establish antimicrobial activity.

### Validation of the chersinamycin BGC using CRISPR-Cas9 gene editing

To confirm that the *M. chersina* BGC identified by genome mining was responsible for chersinamycin production, we performed an LC-MS screen of the mutant strain *M. chersina* ΔPKS7. This strain was constructed in an elegant investigation of the biosynthetic genes responsible for production of the anthraquinone scaffold in dynemicin biosynthesis^66^ and generously gifted to us by the Townsend Lab. It contains a 5.297 kilobase CRISPR-Cas9-mediated knockout of five genes encoding the putative biosynthesis enzymes for Dpg (Chers 29–33, **Figure 6A,B**). We observed that deletion of these biosynthetic genes resulted in the inability of *M. chersina* to produce chersinamycin. We rescued the knockout phenotype by the addition of 1 mM Dpg to the production medium (**Figure 6C**). These studies establish the identity of the chersinamycin BGC and, importantly, demonstrated feasibility of genetic manipulation of this cluster.

**Figure 6.**
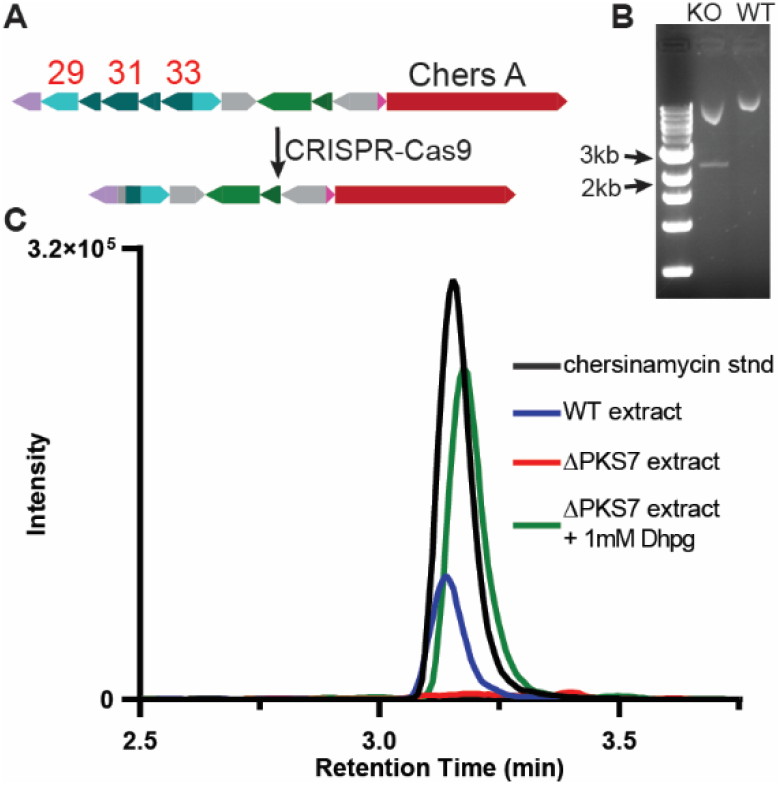
Confirmation of the chersinamycin gene cluster. A) CRISPR-Cas9 facilitated knockout of five genes within the biosynthetic pathway of chersinamycin. The genes have homology to PLP-dependent aminotransferase (Chers 29), DpgD (Chers 30), DpgC (Chers 31), DpgB (Chers 32), and DpgA (Chers 33). B) Confirmation of the knockout region in ΔPKS7 strain visualized by a 2.2 kb band generated from PCR of gDNA with primers flanking the knockout region. C) Extracted ion chromatograms for the doubly charged ion species of chersinamycin (m/z = 1288) in a chersinamycin standard and crude extracts from wild-type *M. chersina*, ΔPKS7, and ΔPKS7 complemented with 1 mM Dpg.

### Assessment of antimicrobial activity of chersinamycin

Chersinamycin was examined for its ability to inhibit bacterial growth by broth microdilution assays against Gram-positive strains *B. subtilis* ATCC 6051, *S. aureus* ATCC 25923, and *E. faecalis* ATCC 29212 and Gram-negative strain *E. coli* ATCC 25922. As our SAR-based search criteria biased our genome mining toward an active compound, we expected antimicrobial activity and mode of action to be consistent with what has been established for this family of antibiotics. Chersinamycin was found to be ineffective against *E. coli* but have potent antimicrobial activity against the Gram-positive strains (**Table 1**). Due to its structural similarities to ramoplanin, we expect activity against important clinically relevant pathogens such as *C. difficile* as well. As such, chersinamycin provides an additional potent ramoplanin family antibiotic for investigation into its antimicrobial potency and pharmacokinetic properties.

**Table 1.** MICs of ramoplanin and chersinamycin as measured by broth microdilution assay.

|  | Ramoplanin | Chersinamycin |
| --- | --- | --- |
| *B. subtilis* ATCC 6051 | < 0.125 µg mL<sup>-1</sup> | < 0.125 µg mL<sup>-1</sup> |
| *S. aureus* ATCC 25923 | 0.5 µg mL<sup>-1</sup> | 2 µg mL<sup>-1</sup> |
| *E. faecalis* ATCC 29212 | 0.5 µg mL<sup>-1</sup> | 1 µg mL<sup>-1</sup> |
| *E. coli* ATCC 25922 | >64 µg mL<sup>-1</sup> | >64 µg mL<sup>-1</sup> |

## CONCLUSIONS

The emergence of resistance to nearly all our first line antibiotics has put enormous pressure on the development of new therapeutics. Ramoplanin is a potent antibiotic that is bactericidal against a number of clinically relevant Gram-positive pathogens, but poor bioavailability and stability highlight a need for development of next generation analogs with better pharmacological properties. We have reported a targeted genome mining strategy able to rapidly and reliably identify ramoplanin family gene clusters using established SAR. This has resulted in the discovery of five previously unidentified ramoplanin family BGCs in five additional bacterial strains. Of the strains identified, four have been previously cultured and extracted for other biologically active natural products, highlighting the importance of precise screening and extraction methods in identifying new natural products, and the significance of genome mining in natural product discovery. Bioinformatic analyses of putative proteins within the gene clusters allowed for structural predictions of the encoded natural products. These analyses predict 17-residue lipoglycodepsipeptides (from *M. chersina* and *A. orientalis* strains) and lipodepsipeptides (from *A. balhimycina* and *Streptomyces* sp. TLI_053) with high sequence similarity to ramoplanin and enduracidin, providing further support of what our group and others had concluded through SAR as to the significance of certain structural features for this class of antibiotics. Bettering our understanding of SAR through such analyses will aid in more insightful design of new antibiotics with improved biological properties.

To validate one of the five identified biosynthetic gene clusters, we isolated the new antibiotic chersinamycin from fermentation of *M. chersina*. We evaluated its covalent structure, and we used a CRISPR-Cas9-generated mutant strain to validate that this gene cluster produces chersinamycin. Thorough bioinformatic analysis paired with classical structure determination approaches allowed for structure elucidation, thus expanding this important antibiotic class for the first time since the discovery of ramoplanin over three decades ago. Chersinamycin retains many of the structural features of ramoplanin, including the presence of two mannose sugars which have been demonstrated to contribute to ramoplanin’s stability and improved solubility over its sister compound enduracidin. The peptide was determined to have a saturated N-acyl lipid, contrasting the lipid structures of the other two characterized compounds within this family and consistent with the lack of dehydrogenases within the identified gene cluster. Interestingly, the gene cluster retains the oxidoreductase (Chers 44) which has been hypothesized to play a role in lipid unsaturation. Therefore, further investigation is needed to understand the lipid biosynthetic pathway in this antibiotic class, greater understanding of which may aid in the development of biosynthetic analogs with new lipid architectures of decreased hemolytic activity.^16^

Finally, the isolation of a ramoplanin family compound from a genetically tractable strain provides exciting opportunities for investigation of the biosynthetic pathway and development of biosynthetic analogs. Independently from this study, a CRISPR-Cas9 strategy was developed to produce a series of gene-inactivation mutants throughout the genome of *M. chersina*, a strategy that is difficult to achieve in many strains of natural product-producing organisms.^66^ We have demonstrated that one such mutant strain, *M. chersina* ΔPKS7, contains a knockout of the Dpg biosynthesis genes within the chersinamycin BGC that abolishes chersinamycin production. Our ability to rescue production through supplementation of Dpg in the production medium demonstrates the feasibility of CRISPR-mediated manipulation of this biosynthetic pathway. This work therefore presents exciting opportunities for targeted gene inactivation to investigate enzymes within the chersinamycin biosynthetic pathway, as well as to produce biosynthetic analogs.

## MATERIALS AND METHODS

### General methods and materials

Bacterial cell culture media components were purchased from Affymetrix, Fisher Scientific, Millipore-Sigma, and BD Difco Laboratories. A sample of Pharmamedia was obtained from Archer Daniels Midland Company, and fish meal was purchased from Coyote Creek Organic Feed Mill and Farm. Ultra-high purity solvents were purchased from Millipore-Sigma and Fisher Scientific and used without further purification. All chemicals were purchased in their highest purity forms from Millipore-Sigma and used without further purification unless otherwise indicated. The 1D and 2D NMR spectra (COSY, TOCSY, NOESY) were collected on a Varian/Agilent DirectDrive2 spectrometer at 800 MHz. Preparative reverse-phase HPLC purifications were performed on a Waters Prep 150B system with a Phenomenex octadecyl silica (C_18_) column (250 mm × 21 mm, 10 μm, 300 Å) or Vydac C_18_ column (250 × 10 mm, 5 μm, 300 Å). Analytical HPLC was performed on a Varian Prostar system with a Phenomenex C_18_ column (250 × 4.6 mm, 5 μm, 300 Å). Tandem MS/MS spectrometry was performed using a Fusion Lumos Orbitrap mass spectrometer. Matrix-assisted laser desorption time-of-flight mass spectrometry (MALDI-TOF) was performed using a Bruker Autoflex Speed LRF MALDI-TOF System. High-resolution mass spectra were collected on an Agilent 6224 LC/MS-TOF instrument.

### Bioinformatics

The NCBI accession numbers for the ramoplanin and enduracidin biosynthetic gene loci are DD382878 and DQ403252, respectively. NRPS A, NRPS B, NRPS C, NRPS D, the terminal thioesterase subdomain from NRPS C, the FAAL, and the ACP were used as initial queries for protein blast searches against the NCBI database. Sequences with >50% identity were collected and organisms that had four or more homologous proteins to the search queries were considered hits. Whole genome sequences for these organisms were obtained from NCBI GenBank and open reading frames within 40 ORFs on either side of NRPS B were analyzed. A total of 1069 translated sequences were subjected to an all vs. all blast and assembled into a sequence similarity network with an E value limit of 10^−5^ and alignment score of 50 using EFI-Enzyme Similarity Tool. The network was visualized using Cytoscape (version 3.7.1, from the National Resource of Network Biology). From the initial network five genomes were selected as having enough clustered proteins for a full BGC and were assembled into a more targeted SSN using an E value limit of 10^−5^ and alignment scores of 25 and 50. Manual analysis was complemented with antiSMASH 4.0 using the following: FMIB01000002.1 (*M. chersina* strain DSM 44151, cluster 1), NZ_CP016174 (*A. orientalis* strain B-37, cluster 13) NZ_ASJB01000042 (*A. orientalis* strain DSM 40040), NZ_KB913037 (*A. balhimycina* FH 1894 strain DSM 44591, clusters 1, 28), NZ_LT629775 (*Streptomyces* sp. TLI_053, cluster 18).

### Antibiotic production screening in *M. chersina* DSM 44151

To prepare the seed culture, a frozen aliquot of *M. chersina* vegetative stock (4 mL, See Supplemental Information) was thawed on ice, then used to inoculate a 500 mL baffled flask containing 100 mL of medium 53 and was incubated at 28 °C for seven days with shaking at 250 rpm. For antibiotic production, seed culture (4 mL) was used to inoculate a 500 mL flask containing 100 mL of each of following media: dynemicin production medium H881 (10 g L^−1^ starch; 5 g L^−1^ Pharmamedia; 1 g L^−1^ CaCO_3_; 0.05 g L^−1^ CuSO_4_; and 0.5 mg L^−1^ NaI);^62^ H881 medium with chicken oil (14 mL L^−1^); H881 medium with glucose (30 g L^−1^); enduracidin growth medium (80 g L^−1^ corn flour; 30 g L^−1^ corn gluten meal; 5 mL L^−1^ corn steep liquor; 3 g L^−1^ ammonium sulfate; 1 g L^−1^ NaCl; 10 mg L^−1^ ZnCl_2_; 10 g L^−1^ lactose; 10 mL L^−1^ potassium lactate; and 14 mL L^−1^ chicken oil),^61^ or ramoplanin production medium (50 g L^−1^ starch; 30 g L^−1^ glucose; 30 g L^−1^ soy flour; 10 g L^−1^ CaCO_3_; 5 g L^−1^ leucine).^39^ The chicken oil supplement was prepared by defatting 1 whole roasting chicken (Harris Teeter, Inc.), rendering the isolated fat and skin at 350 °C for 15 min, cooling the mixture to rt, and clarifying the oil by centrifugation (15 min, 4,000 rpm, 4 °C). The oil was stored in the dark at 4 °C for up to 2 days prior to use.

Production cultures of *M. chersina* were grown at 28 °C, 250 rpm for 12–21 days. Antibiotic production was monitored by MALDI-TOF MS screening. For screening, cell culture aliquots (6 mL) were pelleted by centrifugation at 5000 rpm for 15 minutes at 4 °C. The supernatant was separated from the cell pellet by decantation and the supernatant fraction was extracted with ethyl acetate. The organic fraction was separated, dried with sodium sulfate, and freed of solvent under vacuum. Both the aqueous and organic fractions were analyzed by MALDI-TOF MS for production of secondary metabolites in the 2000–3000 Da MW range. Similarly, the production culture aliquot cell pellet was resuspended in acidic aqueous MeOH/H_2_O (66:33 v/v; pH 3, 6 mL), stirred at rt for 3 h to affect cell lysis, centrifuged (5000 rpm, 10 min, 4 °C), and the supernatant was decanted and extracted with EtOAc as above. Both the aqueous and organic fractions were analyzed by MALDI-TOF MS. The antibiotic peptide was observed in the aqueous fraction of the extracted cell pellet, which was used for further analyses.

### Large scale production, isolation, and purification of chersinamycin from *M. chersina* DSM 44151

For large scale production of chersinamycin from *M. chersina*, 20 mL of seed culture was used to inoculate 2 L baffled flasks containing 500 mL H881 media and grown at 28 °C, 250 rpm for 12 days. Cells were pelleted by centrifugation, resuspended in acidic aqueous MeOH (300 mL), stirred at rt for 3 h at rt, then centrifuged to remove cellular debris as described above. The supernatant was extracted with EtOAc (3 × 300 mL) to remove organic-soluble metabolites. The aqueous layer was freeze-dried, dissolved in an H_2_O/MeCN mixture, and subjected to RP-HPLC using a Jupiter C18, 250 × 21.2 mm column with a linear gradient of 20–50% B over 30 minutes, where solvent A was 0.1% TFA in H_2_O and B was 0.06% TFA in MeCN. A second HPLC purification was performed using a Vydac C18 250 × 10 mm column with the same solvent system as above and a linear gradient of 20–35% B over 50 minutes to yield pure chersinamycin in 1–3 mg L^−1^ quantities from the starting cell culture.

### Advanced Marfey’s analysis of chersinamycin and ramoplanin

To facilitate the hydrolysis of chersinamycin and ramoplanin for advanced Marfey’s analysis, to a thick walled glass vial (10 mL) containing either lyophilized chersinamycin (0.8 mg, 311 pmol) or ramoplanin (1 mg, 392 pmol) was added freshly prepared 6 M HCl (200 μL). After flushing the vial with Ar for 20 min, the vial was sealed and heated at 110 °C for 18 hrs. The reaction mixtures were cooled, evaporated under a stream of N_2_, dissolved in TEA/H_2_O (25:75, v/v, 100 μL), transferred to a 5 mL round bottom flask, and evaporated under reduced pressure to dryness. The latter sequence was repeated 2 additional times. The resulting residue was dissolved in H_2_O (75 μL), sodium bicarbonate (1M, 40 μL) and TEA (25 μL) were added, and the mixture was transferred to a 1.7 mL amber Eppendorf tube. Marfey’s reagent (1.4 mg) in acetone (100 μL) was added and the mixture was heated for 1h at 40 °C with periodic vortexing. After cooling to rt, HCl (2M, 10 μL) was added and the reaction mixture was dried overnight in a vacuum desiccator. For HPLC analysis, dried reaction mixtures were dissolved in DMSO (0.5 mL). A 50 μL aliquot was used to make a 1:1 dilution in water and filtered through a 0.2 μm syringe filter. RP-HPLC-MS analysis was performed with at Kintex 2.6 μm EVO-C18, 100 × 3 mm column with a gradient of 5–50% B over 40 minutes, where solvent A was 100:3:0.3 H_2_O/MeOH/TFA and solvent B was 100:3:0.3 MeCN/H_2_O/TFA. ESI-MS for FDAA-amino acids was performed in negative ion mode.

### Structural determination by 1D and 2D NMR and ESI-MS/MS

Pure chersinamycin (3 mg, 2.6 mM) was dissolved in 4:1 H_2_O/DMSO-*d_6_* (v/v) or 4:1 D_2_O/DMSO-*d_6_* at pH 4.56. Homonuclear experiments were acquired with a spectral width of 11 ppm. Mixing times of 80 and 500 ms were used for TOCSY and NOESY spectra, respectively. Solvent suppression was employed at 2.50 ppm (DMSO) and 4.54 ppm (H_2_O) and spectra were referenced to DMSO. For ESI-MS/MS analysis, pure cyclic and acyclic peptides dissolved in 4:1 H_2_O/MeCN (v/v) were diluted 1:20 with 1:1 H_2_O/MeCN (v/v) with 0.2% formic acid and infused into a Fusion Lumos Orbitrap mass spectrometer at 2.5 μL min^−1^. Data was collected at 120 K for full MS scans and 30 K for MS/MS scans. The intact peptide was subjected to MS/MS higher-energy C-trap dissociation (HCD) fragmentation in both the [M+2H]^2+^ and [M+3H]^3+^ charge states.

### Genetic and biochemical confirmation of antibiotic production by the predicted chersinamycin BGC

A frozen aliquot (100 μL) of mycelia from the *M. chersina* Dpg deletion mutant strain ΔPKS7^66^ was thawed on ice, plated onto medium 53 agar, and incubated at 28 °C for five days. Sterile liquid medium 53 was added to the plate (2 mL) and the plate was scraped to resuspend the cells. This suspension was added to a sterile culture flask (125 mL) containing medium 53 (50 mL), and the mixture was incubated for seven days at 28 °C with shaking at 250 rpm. An aliquot of this seed culture (2 mL) was used to inoculate H881 media (50 mL) in a 250 mL sterile culture flask, which was incubated at 28 °C for 12 days with shaking (250 rpm). Following centrifugation, the production cell pellet was extracted with acidic aqueous MeOH/H_2_O (66:33 v/v; pH 3, 50 mL) for 3 hours at rt. Cell debris was removed by centrifugation and the supernatant was subjected to HPLC-MS analysis for validation of the absence of detectible chersinamycin. To restore chersinamycin production through chemical complementation, *M. chersina* strain ΔPKS7 was fermented in H881 production medium that was supplemented with racemic (*R*,*S*)-3,5-Dpg (1 mM, Millipore-Sigma). Production cultures were incubated identically as above, the cell pellets were isolated by centrifugation, and then extracted and analyzed by HPLC-MS.

### Minimal inhibitory concentration assays

Antibacterial activity of chersinamycin and positive controls (vancomycin, ampicillin, and ramoplanin A2,) were determined by the broth microdilution assay method. Briefly, bacterial strains were grown in cation-adjusted Mueller–Hinton broth. A microtiter plate was prepared by coating wells in 0.2% BSA, and antimicrobial peptides were added with 2-fold dilution steps ranging from 64–0.125 μg mL^−1^. Bacteria was added to a final concentration of 10^5^ colony forming units and final volume of 100 μL. Plates were incubated at 37 °C for 24 hours, and the MIC was read as the lowest peptide concentration for which no bacterial growth was visualized. Reported values are the average of two replicates.

## ASSOCIATED CONTENT

### Supporting Information

The Supporting Information is available and contains:

General methods of bacterial culture, antibiotic screening in *A. orientalis* and *A. balhimycina,* and semisynthetic derivatizations of chersinamycin.
Detailed annotations of the bacterial strains and biosynthetic gene clusters analyzed in this study, MS and NMR data for chersinamycin

### Accession Codes

Ramoplanin biosynthetic gene cluster, Accession DD382878; Enduracidin biosynthetic gene cluster, DQ403252; *Micromonospora chersina* DSM 44151, Accession FMIB01000002.1; *Amycolatopsis orientalis* strain B-37, Accession NZ_CP016174; *Amycolatopsis orientalis* DSM 40040 = KCTC 4912, Accession NZ_ASJB01000042; *Amycolatopsis balhimycina* FH 1894 DSM 44591, Accession NZ_KB913037; *Streptomyces* sp. TLI_053, Accession NZ_LT629775; *Micromonospora* sp. MH33, Accession NZ_MUYZ00000000.1; *Amycolatopsis thailandensis* strain JCM 16380, Accession NZ_NMQT00000000.1; *Actinomadura madurae* LIID-AJ290, Accession NZ_AWOO02000001.1; *Actinomadura madurae* strain DSM 43067, Accession NZ_FOVH00000000.1; *Streptomyces vietnamensis* strain GIM4.0001, Accession NZ_CP010407.1; *Streptomyces* sp. GP55, Accession NZ_PJMT01000001.1; *Streptomyces cinnamoneus* strain ATCC 21532, Accession NZ_NHZO00000000.1; *Streptomyces cinnamoneus* strain DSM 41675, Accession NZ_PKFQ01000001.1

## AUTHOR INFORMATION

### Notes

The authors declare no competing financial interests.

## ACKNOWLEDGMENTS

The authors gratefully acknowledge Drs. Craig Townsend and Douglas Cohen for the generous gift of *Micromonospora chersina* strain ΔPKS7, Dr. Peter Silinski for mass spectrometry assistance, Drs. Anthony Ribeiro, Benjamin Bobay, and the Duke NMR Facility for NMR assistance, and the Duke University School of Medicine for the use of Proteomics and Metabolomics Shared Resource, which provided MS/MS sequencing of chersinamycin.

## ABBREVIATIONS

PG: peptidoglycan
BGC: biosynthetic gene cluster
NRPS: nonribosomal peptide synthetase
ORF: open reading frame
FAAL: fatty acyl-AMP ligase
ACP: acyl carrier protein
ACAD: acyl-CoA dehydrogenase
SSN: sequence similarity network
Hpg: 4-hydroxyphenylglycine
Dpg: 3,5-dihydroxyphenylglycine
Orn: ornithine
End: enduracididine
ClHpg: 3-chlorohydroxyphenylglycine
diClHpg: 3,5-dichlorohydroxyphenylglycine

