## Supplemental Information for "Targeted Genome Mining Discovery of the Ramoplanin Congener Chersinamycin from the Dynemicin-Producer *Micromonospora chersina* DSM 44154"

**Contents:**

**Supplementary Methods**

**List of Tables**

**Table S1.**	Identified bacterial strains with homologs to key ramoplanin and enduracidin biosynthesis proteins**.**

**Table S2.**	Comparison of the ramoplanin-family gene clusters in seven Actinobacteria strains.

**Table S3.**	Amino acid sequence comparison of predicted peptide products from ramoplanin family BGCs.

**Table S4.**	NMR spectroscopic data of chersinamycin.

**Table S5.**	List of calculated and observed b- and y-ions from MS/MS of acyclic chersinamycin**.**

**Table S6.**	Retention time for FDAA derivatives of amino acid standards and chersinamycin hydrolysate.

**Table S7.**	Deduced functions of proteins within the defined BGC of *M. chersina* strain DSM 44151.

**Table S8.**	Deduced functions of proteins within the defined BGC of *A. orientalis* strain B-37.

**Table S9.**	Deduced functions of proteins within the defined BGC of *A. orientalis* strain DSM 40040.

**Table S10.**	Deduced functions of proteins within the defined BGC of *A. balhimycina* FH 1894 strain DSM 44591.

**Table S11.**	Deduced functions of proteins within the defined BGC of *Streptomyces* sp. TLI_053.

**List of Figures**

**Figure S1.**	Sequence similarity network of ORFs surrounding NRPS proteins in new bacterial strains.

**Figure S2.**	Open reading frame and NRPS domain comparisons between ramoplanin family gene clusters.

**Figure S3.**	Phylogenetic relationship between NRPS condensation domains.

**Figure S4.**	Phylogenetic relationship between terminal NRPS thioesterase domains.

**Figure S5.**	HR-ESI-MS of chersinamycin.

**Figure S6.**	HR-ESI-MS of acyclic chersinamycin.

**Figure S7.**	ESI-MS spectrum of di-propionylated chersinamycin

**Figure S8.**	MALDI-MS spectrum of hydrogenated ramoplanin and chersinamycin

**Figure S9.**	ESI-MS/MS spectrum of chersinamycin.

**Figure S10.**	ESI-MS/MS spectrum of acyclic chersinamycin.

**Figure S11.**	MS/MS fragmentation of acyclic chersinamycin (b- and y-ion series).

**Figure S12.**	Determination of absolute configuration of amino acids by advanced Marfey’s analysis.

**Figure S13.**	^1^H NMR (800 MHz, 4:1 H_2_O/DMSO-*d_6_*) spectrum of chersinamycin.

**Figure S14.**	^1^H-^1^H COSY (800 MHz, 4:1 H_2_O/DMSO-*d_6_*) spectrum of chersinamycin

**Figure S15.**	^1^H-^1^H TOCSY (800 MHz, 4:1 H_2_O/DMSO-*d_6_*) spectrum of chersinamycin

**Figure S16.**	^1^H-^1^H NOESY (800 MHz, 4:1 H_2_O/DMSO-*d_6_*) spectrum of chersinamycin

**Figure S17.**	^1^H-^1^H NOESY (800 MHz, DMSO-*d_6_*) spectrum of chersinamycin

**Figure S18.**	Depiction of defining NMR correlations observed in chersinamycin.

**Bacterial strains and culture conditions**. *Micromonospora chersina* strain DSM 44151 was purchased from the ATCC and cultivated as reported by Lam et al.^1^ Briefly, freeze-dried *M. chersina* was reconstituted and grown on ISP 2 agar plates at 26 °C for 4 days until spore formation was visible. Spores were collected according to established protocols^2^ and used to inoculate 100 mL of seed medium 53 (10 g L^-1^ fish meal; 30 g L^-1^ dextrin; 10 g L^-1^: lactose; 6 g L^-1^ CaSO_4_; and 5 g L^-1^ CaCO_3_) in a 250 mL culture flask, which was incubated for 7 days at 28 °C with orbital agitation at 250 rpm. Frozen vegetative stocks of *M. chersina* were prepared by mixing the seed culture suspension with an equal volume of 20% glycerol/10% sucrose, which was subsequently aliquoted, flash frozen with liquid nitrogen, and stored at -80 °C.

*Amycolatopsis orientalis* strain DSM 40040 was purchased from the Leibniz Institute DSMZ. Freeze-dried *A. orientalis* was reconstituted in ISP I medium and plated onto ISP II agar plates. Plates were incubated at 26 °C for 5 days, after which the lawn of bacteria was lifted by adding sterile water (1 mL) and scraping gently with a sterile cell spreader. The suspension was used to inoculate 40 mL of vancomycin seed medium (5 g L^-1^ glucose; 10 g L^-1^ starch; 5 g L^-1^ peptone; and 2 g L^-1^ yeast extract)^3^ in a 250 mL culture flask, which was incubated for 2 days at 30 °C with orbital agitation at 220 rpm. Frozen vegetative stocks were prepared by mixing the seed culture suspension with an equal volume of 80% glycerol, which was subsequently aliquoted, flash frozen in liquid nitrogen, and stored at -80 °C.

*Amycolatopsis balhimycina* FH 1894 strain DSM 44591 was purchased from the Leibniz Institute DSMZ. Freeze-dried *A. balhimycina* was reconstituted in GYM Streptomyces liquid medium and plated onto GYM Streptomyces agar plates. Agar plates were incubated at 28 °C for 4 days, after which the lawn of bacteria was lifted by adding sterile water (1 mL) and scraping gently with a sterile cell spreader. The suspension was used to inoculate 25 mL of tryptic soy broth in a 125 mL culture flask, which was incubated for 2 days at 28 °C with orbital agitation at 220 rpm. Frozen vegetative stocks were prepared by mixing culture suspension with an equal volume of 80% glycerol, which was subsequently aliquoted, flash frozen in liquid nitrogen, and stored at -80 °C.

**Macrolactone selective hydrolysis**. Triethylamine (3 μL) was added to chersinamycin dissolved in water (0.115 μmol, 297 μL) to give 1% (v/v) TEA. The solution was allowed to sit at room temperature for one hour, and then analyzed by MALDI-TOF. After determining that the reaction had gone to completion by complete consumption of the starting material, the reaction mixture was dried and reconstituted in a water/acetonitrile mixture for further MS/MS analyses. Acyclic chersinamycin ESI-MS (*m/z*): [M+2H]^2+^ calcd for C_119_H_160_ClN_21_O_42_, 1296.044; found, 1296.044

**Catalytic hydrogenation of the N-acyl lipid**. The procedure for catalytic hydrogenation of the N-acyl lipid was modified from that described by Ciabatti and Cavalleri.^4^ Briefly, to a glass conical microvial charged with either ramoplanin A2 or chersinamycin (2 mg), MeOH/H_2_O (10:90, v/v, 389 μL) was added and the solution was stirred at rt to facilitate dissolution. Once dissolved, Pd/C (2.5% w/w) was added (1 mg, 5.0 mol %), the flask was evacuated under vacuum, flushed with argon, and then the reaction mixture was placed under an atmosphere of H_2_ and stirred and monitored by analytical HPLC. After 8h, additional Pd/C (2.5%, 1 mg) was added and the mixture stirred overnight under an H_2_ atmosphere. The reactions were diluted with MeOH/H_2_O (10:90, v/v, 389 μL), filtered through Celite^TM^, dried under vacuum, and analyzed by MALDI-TOF. A mass shift indicated a change from ramoplanin A2 (MALDI-TOF MH 2553.500) to tetrahydroramoplanin A2 (MALDI-TOF MH 2557.731). No mass shift was observed for chersinamycin (MALDI-TOF MH 2573.404).

**Propionic anhydride labeling of chersinamycin**

Propionic anhydride derivatization of chersinamycin was employed to confirm the two predicted ornithine residues and provide a mass spectrometry handle. The derivatization procedure was adapted from the method described by Garcia et al. for histone labeling.^5^ Lyophilized peptide was dissolved in water to a concentration of 1 μg/μL. The peptide (5 μL) was transferred to an Eppendorf tube to which ammonium bicarbonate pH 8 (5 μL, 100 mM) and concentrated ammonium hydroxide (2 μL) were added. Propionic anhydride/methanol (3:1 v/v, 20 μL) was then added. The mixture was briefly vortexed, adjusted to pH 8 with ammonium hydroxide, and incubated at 51 °C for 20 minutes. The mixture was concentrated to 5 μL using a SpeedVac concentrator. To the mixture was added ammonium bicarbonate (5 μL, 100 mM) and the method was repeated. Concentrated samples were diluted in water and analyzed by HR-MS. The observed mass in consistent with the incorporation of to propionyl groups to the core peptide.

Di-propionylated chersinamycin ESI-MS (*m/z*): [M+2H]^2+^ calcd for C_125_H_168_ClN_21_O_43_^2+^, 1343.57; found, 1343.568

Table S1. Identified bacterial strains with homologs to key ramoplanin and enduracidin biosynthesis proteins.

| **Organism/Name** | **NRPS A** | **NRPS B** | **NRPS C** | **NRPS D** | **FAAL** | **ACP** | **Thioesterase** |
| --- | --- | --- | --- | --- | --- | --- | --- |
| *Streptomyces fungicidicus* ATCC 21013 (enduracidin) | R | R | R | R | R | R | R |
| *Micromonospora chersina* strain DSM 44151 | R | R, E | R, E | R, E | R, E | R, E | R, E |
| *Amycolatopsis orientalis* strain B-37 | R, E | R, E | R, E | R, E | R, E | R, E | R, E |
| *Amycolatopsis orientalis* DSM 40040 = KCTC 9412 | R, E | R, E | R, E | R, E | R, E | R, E | R, E |
| *Amycolatopsis balhimycina* FH 1894 strain DSM 44591 | R, E | R, E | R, E | R, E | R, E | R, E | R, E |
| *Streptomyces* sp. TLI_053 |  | R, E | R, E | R, E | R, E | R, E | R, E |
| *Micromonospora* sp. MH33 |  | R, E | R, E | R, E | R, E | R, E | R, E |
| *Amycolatopsis thailandensis* srain JCM 16389 | R, E | R, E | E |  | R, E | R, E | R, E |
| *Actinomadura madurae* LIID-AJ290 |  | R | R, E |  | R, E | R | E |
| *Actinomadura madurae* strain DSM 43067 |  | R | R, E |  | R, E | R | E |
| *Streptomyces vietnamensis* strain GIM4.0001 | E | E |  |  | R, E | R, E |  |
| *Streptomyces* sp. GP55 | E | E |  |  | R, E | R, E |  |
| *Streptomyces cinnamoneus* strain ATCC 21532 |  | R, E |  |  | R, E | R, E | R, E |
| *Streptomyces cinnamoneu*s strain DSM 41675 |  |  | R, E |  | R, E | R, E | R, E |

Analyzed proteins are Ramo A/End A, Ramo B/End B, Ramo C/End C, Ramo D/End D, and each respective FAAL, ACP, and terminal thioesterase of NRPS C. An R indicates >50% identity to the ramoplanin homologue and E indicates >50% identity to the enduracidin homolog.

Table S2. Comparison of the ramoplanin-family gene clusters in seven bacterial strains.

|  | **Enduracidin** | **Ramoplanin** | ***M. chersina*** | ***A. orientalis* B-37** | ***A. orientalis* DSM 40040** | ***A. bahlimycina*** | ***Streptomyces* sp. TLI_053** |
| --- | --- | --- | --- | --- | --- | --- | --- |
| Acetyl-CoA acetyltransferase (thiolase) | Orf 11 |  |  |  |  | Orf 12 43%^a^ |  |
| Transcriptional regulator | Orf 12 |  |  |  |  |  |  |
| -Mannosidase | Orf 13 |  |  |  |  |  |  |
| Probable sugar transport system lipoprotein | Orf 14 |  |  |  |  |  |  |
| Sugar transport system permease protein | Orf 15 |  |  |  |  |  |  |
| Sugar transport system permease protein | Orf 16 |  |  |  |  |  |  |
| Ribonuclease D | Orf 17 |  |  |  |  |  |  |
| Two-component response regulator | Orf 18 |  |  |  |  |  |  |
| Unknown | Orf 19 |  |  |  |  |  |  |
| Uroporphyrinogen decarboxylase | Orf 20 |  |  |  |  |  |  |
| PAS protein phosphatase 2C-like | Orf 21 |  |  |  |  |  |  |
| Str-like regulatory protein | Orf 22 43%^b^ | Orf 5 43%^a^ | Orf 28 44%^a^ 72%^b^ | Orf 29 54%^a^ 45%^b^, Orf 30 44%^a^ 46%^b^ | Orf 31 43%^a^ 47%^b^, Orf 32 54%^a^ 46%^b^ | Orf 29 53%^a^ 46%^b^, Orf 30 45%^a^ 47%^b^ | Orf 36 55%^a^ 41%^b^ |
| Prephenate dehydrogenase | Orf 23 51%^b^ | Orf 4 51%^a^ |  |  |  |  | Orf 37 48%^a^ 52%^b^, Orf 77 57%^a^ 55%^b^ |
| Transcriptional regulator | Orf 24 50%^b^ | Orf 5 49%^a^ | Orf 28 47%^a^ 72%^b^ | Orf 29 49%^a^ 45%^b^, Orf 30 71%^a^ 46%^b^ | Orf 31 70%^a^ 47%^b^, Orf 32 49%^a^ 46%^b^ | Orf 29 50%^a^ 46%^b^, Orf 30 74%^a^ 47%^a^ | Orf 36 46%^a^ 41%^b^ |
| 4-Hydroxyphenylpyruvate dioxygenase (HmaS homologue) | Orf 25 48%^b^ | Orf 30 48%^a^ | Orf 34 41%^a^ 41%^b^ | Orf 31 79%^a^ 49%^b^ | Orf 30 80%^a^ 49%^b^ | Orf 31 78% 48%^b^ | Orf 54 42% 41%^b^ |
| Unknown (MppR homologue) | Orf 26 |  |  |  |  |  |  |
| PLP-dependent aminotransferase (MppQ homologue) | Orf 27 |  |  |  |  |  |  |
| PLP-dependent aminotransferase (MppP homologue) | Orf 28 |  |  |  |  |  |  |
| Aminotransferase | Orf 29 | Orf 6 68%^a^, Orf 7 70%^a^ | Orf 60 59%^a^, Orf 29 70%^a^ | Orf 32 78%^a^ | Orf 29 78%^a^ | Orf 32 79%^a^ | Orf 53 67%^a^ |
| FAD-dependent oxidoreductase (halogenase) | Orf 30 64%^b^ | Orf 20 64%^a^ | Orf 49 63%^a^ 83%^b^ | Orf 34 83%^a^ 64%^b^ | Orf 27 83%^a^ 64%^b^ | Orf 34 84%^a^ 64%^b^ |  |
| Transmembrane transport protein | Orf 31 | Orf 1 50%^a^, Orf 3 56^a^ |  | Orf 35 71%^a^ | Orf 26 72%^a^ | Orf 35 73%^a^ | Orf 57 43%^a^ |
| ABC transporter ATP-binding protein | Orf 32 | Orf 23 56%^a^, Orf 2 71%^a^ | Orf 53 73%^a^ | Orf 36 78%^a^ | Orf 25 78%^a^ | Orf 36 81%^a^ | Orf 58 64%^a^ |
| ABC transporter | Orf 33 73%^b^ | Orf 8 73%^a^ | Orf 36 78%^a^ 77%^b^ | Orf 37 78%^a^ 74%^b^ | Orf 24 78%^a^ 74%^b^ | Orf 37 79%^a^ 75%^b^ | Orf 56 62%^a^ 63%^b^ |
| Alpha/beta fold hydrolase | Orf 34 77%^b^ | Orf 9 77%^a^ | Orf 37 75%^a^ 78%^b^ | Orf 38 71%^a^ 73%^b^ | Orf 23 69%^a^ 72%^b^ | Orf 38 76%^a^ 77%^b^ | Orf 55 62%^a^ 63%^b^ |
| MBL fold metallo-hydrolase |  | Orf 10 | Orf 38 82%^b^ |  |  |  | Orf 48 72%^b^ |
| Acyl carrier protein | Orf 35 69%^b^ | Orf 11 69%^a^ | Orf 39 63%^a^ 58%^b^ | Orf 39 75%^a^ 67%^b^ | Orf 22 76%^a^ 66%^b^ | Orf 39 78%^a^ 71%^b^ | Orf 43 54%^a^ 61%^b^ |
| **NRPS A** | **End A 55%^b^** | **Ramo A 55%^a^** | **Orf 40 47%^a^ 61%^b^** | **Orf 40 67%^a^ 54%^b^** | **Orf 21 66%^a^ 53%^b^** | **Orf 40 66%^a^ 55%^b^** | **Orf 42 44%^a^ 48%^b^** |
| **NRPS B** | **End B 62%^b^** | **Ramo B 62%^a^** | **Orf 41 68%^a^ 67%^b^** | **Orf 41 70%^a^ 61%^b^** | **Orf 20 70%^a^ 61%^b^** | **Orf 41a 72%^a^ 66%^b^, Orf 41b 64%^a^ 64%^b^** | **Orf 41 62%^a^ 60%^b^** |
| **NRPS C** | **End C 61%^b^** | **Ramo C 61%^a^** | **Orf 42 64%^a^ 65%^b^** | **Orf 42 71%^a^ 61%^b^** | **Orf 19 71%^a^ 61%^b^** | **Orf 42 72%^a^ 61%^b^** | **Orf 40 62%^a^ 60%^b^** |
| Thioesterase | EndC 66%^b^ | Orf 15 66%^a^ | Orf 43 70%^a^ 70%^b^ | Orf 43) 79%^a^ 64%^b^ | Orf 18 79%^a^ 65%^b^ | Orf 43 83%^a^ 64%^b^ | Orf 64 55%^a^ 53%^b^ |
| NAD(P)-dependent oxidoreductase | Orf 39 80%^b^ | Orf 16 80%^a^ | Orf 44 81%^a^ 84%^b^ | Orf 44 85%^a^ 78%^b^ | Orf 17 85%^a^ 79%^b^ | Orf 44 86%^a^ 78%^b^ | Orf 63 69%^a^ 71%^b^ |
| **NRPS D** | **End D 57%^b^** | **Ramo D 57%^a^** | **Orf 45 63%^a^ 63%^a^** | **Orf 45 67%^a^ 58%^b^** | **Orf 16 67%^a^ 57%^b^** | **Orf 45 69%^a^ 59%^b^** | **Orf 62 46%^a^ 46%^b^** |
| Hypothetical protein GA0070603_0076 |  | Orf 18 | Orf 47 48%^b^ |  |  |  |  |
| DUF2029 domain-containing protein |  | Orf 19 | Orf 48 68%^b^ |  |  |  |  |
| DNA-binding response regulator | Orf 41 71%^b^ | Orf 21 71%^a^ | Orf 50 76%^a^ 82%^b^ | Orf 46 74%^a^ 70%^b^ | Orf 15 75%^a^ 71%^b^ | Orf 46 77%^a^ 73%^b^ | Orf 61 70%^a^ 70%^b^ |
| Sensor histidine kinase | Orf 42 57%^b^ | Orf 22 57%^a^ | Orf 51 63%^a^ 61%^b^ | Orf 47 72%^a^ 55%^b^ | Orf 14 72%^a^ 55%^b^ | Orf 47 74%^a^ 56%^b^ |  |
| Two-component sensor histidine kinase | Orf 43 |  |  | Orf 48 56%^a^ | Orf 13 56%^a^ | Orf 48 55%^a^ |  |
| Acyl-coA dehydrogenase | Orf 44 67%^b^ | Orf 24 67%^a^ |  | Orf 50 79%^a^ 57%^b^ | Orf 11 78%^a^ 66%^b^ | Orf 49 78%^a^ 65%^b^ | Orf 44 69%^a^ 67%^b^ |
| Acyl-CoA ligase (FAAL) | Orf 45 54%^b^ | Orf 26 54%^a^ | Orf 54 62%^a^ 63%^b^ | Orf 52 69%^a^ 59%^b^ | Orf 9 69%^a^ 59%^b^ | Orf 51 69%^a^ 59%^b^ | Orf 46 51%^a^ 54%^b^ |
| Acyl-CoA dehydrogenase | Orf 45 64%^b^ | Orf 25 64%^a^ |  | Orf 51 74%^a^ 65%^b^ | Orf 10 74%^a^ 65%^b^ | Orf 50 78%^a^ 65%^b^ | Orf 45 69%^a^ 64%^b^ |
| MbtH-like protein | Orf 46 89%^b^ | Orf 27 89%^a^ | Orf 55 91%^a^ 93%^b^ | Orf 53 90%^a^ 87%^b^ | Orf 8 90%^a^ 87%^b^ | Orf 52 91%^a^ 88%^b^ | Orf 47) 82%^a^ 82%^b^ |
| Chorismate mutase |  | Orf 28 | Orf 58 65%^b^ |  |  |  |  |
| Glycosyltransferase |  | Orf 29 | Orf 59 59%^b^ | Orf 49 55%^b^ | Orf 12 64%^b^ |  |  |
| Integral membrane protein | Orf 47 |  |  |  |  |  |  |
| Integral membrane protein | Orf 48 |  |  |  |  |  |  |
| Putative membrane antiporter |  | Orf 31 | Orf 57 34%^b^ |  |  |  |  |

Percent identities are shown for proteins encoded by each ORF compared to the ^a^enduracidin BGC and ^b^ramoplanin BGCs. NRPSs are bolded.

Table S3. Amino acid sequence comparison of predicted peptide products from ramoplanin family BGCs.

| **Module** | **Substrate Recognition Sequence** | **AntiSMASH/NRPSPredictor2** | **Confirmed amino acid** |
| --- | --- | --- | --- |
| NRPS 1 m1 |  |  |  |
| RamoA-m1 | DLTKVGEV | l-Asn | Lipo-l-Asn^1^ |
| EndA-m1 | DLTKVGHV | l-Asp | Lipo-l-Asp^1^ |
| ChersA-m1 | DLTKVGEV | d-Asn | Lipo-Asn^1^ |
| *A. orientalis* B-37-m1 | DLTKVGEV | l-Asn |  |
| *A. orientalis* DSM 40040-m1 | DLTKVGEV | l-Asn |  |
| *A. balhimycina*-m1 | DLTKVGEV | l-Asn |  |
| *Streptomyces* sp. TLI-053-m1 | DLTKVGHI | d-Asp |  |
| NRPS 1 m2 |  |  |  |
| RamoA-m2 | - | - | β-OH-l-Asn^2^ |
| EndA-m2 | DFWSVGMV | l-Thr | l-Thr^2^ |
| ChersA-m2 | DLTKVGEV | l-Asn | β-OH-l-Asn^2^ |
| *A. orientalis* B-37-m2 | DFWSVGMV | l-Thr |  |
| *A. orientalis* DSM 40040-m2 | DFWSVGMV | l-Thr |  |
| *A. balhimycina*-m2 | DFWSVGMV | l-Thr |  |
| *Streptomyces* sp. TLI-053-m2 | DLTKVGHI | l-Asp |  |
| NRPS 2 m1 |  |  |  |
| RamoB-m1 | DAYHLGLL | d-Hpg | d-Hpg^3^ |
| EndB-m1 | DAYHLGLL | d-Hpg | d-Hpg^3^ |
| ChersB-m1 | DAYHLGLL | d-Hpg | d-Hpg^3^ |
| *A. orientalis* B-37-m1 | DAYALGLL | d-Hpg |  |
| *A. orientalis* DSM 40040-m1 | DAYHLGLL | d-Hpg |  |
| *A. balhimycina*-m1 | No sequencing data |  |  |
| *Streptomyces* sp. TLI-053-m1 | DAYHLGLL | d-Hpg |  |
| NRPS 2 m2 |  |  |  |
| RamoB-m2 | DMDTLVSV | d-Tyr, Bht | d-Orn^4^ |
| EndB-m2 | DMETDGSV | d-Orn, Lys, Arg | d-Orn^4^ |
| ChersB-m2 | DMETDGSV | d-Orn, Lys, Arg | d-Orn^4^ |
| *A. orientalis* B-37-m2 | DMET-GSV | d-Orn, Lys, Arg |  |
| *A. orientalis* DSM 40040-m2 | DMETDGSV | d-Orn, Lys, Arg |  |
| *A. balhimycina*-m2 | No sequencing data |  |  |
| *Streptomyces* sp. TLI-053-m2 | DVWHFGQI | d-Glu |  |
| NRPS 2 m3 |  |  |  |
| RamoB-m3 | DFWSVGMW | d-Thr | d-*allo*-Thr^5^ |
| EndB-m3 | DFWSVGMV | d-Thr | d-*allo*-Thr^5^ |
| ChersB-m3 | DFWSVGMV | d-Thr | d-*allo*-Thr^5^ |
| *A. orientalis* B-37-m3 | DLES-GTV | d-Orn, Lys, Arg |  |
| *A. orientalis* DSM 40040-m3 | DLESDGTV | d-Orn, Lys, Arg |  |
| *A. balhimycina*-m3 | No sequencing data |  |  |
| *Streptomyces* sp. TLI-053-m3 | DMETLVSV | d-Orn, Lys, Arg |  |
| NRPS 2 m4 |  |  |  |
| RamoB-m4 | DAYHLGLL | l-Hpg | l-Hpg^6^ |
| EndB-m4 | DAYHLGLL | l-Hpg | l-Hpg^6^ |
| ChersB-m4 | DAYHLGLL | l-Hpg | l-Hpg^6^ |
| *A. orientalis* B-37-m4 | DAYHLGLL | l-Hpg |  |
| *A. orientalis* DSM 40040-m3 | DAYHLGLL | l-Hpg |  |
| *A. balhimycina*-m4 | No sequencing data |  |  |
| *Streptomyces* sp. TLI-053-m4 | DAYHLGLL | l-Hpg |  |
| NRPS 2 m5 |  |  |  |
| RamoB-m5 | DAYHLGLL | d-Hpg | d-Hpg^7^ |
| EndB-m5 | DAYHLGLL | d-Hpg | d-Hpg^7^ |
| ChersB-m5 | DAYHLGLL | d-Hpg | d-Hpg^7^ |
| *A. orientalis* B-37-m5 | DAYALGLL | d-Hpg |  |
| *A. orientalis* DSM 40040-m5 | DAYHLGLL | d-Hpg |  |
| *A. balhimycina*-m5 | No sequencing data |  |  |
| *Streptomyces* sp. TLI-053-m5 | DAYALGLL | d-Hpg |  |
| NRPS 2 m6 |  |  |  |
| RamoB-m6 | No A domain | - | l-*allo*-Thr^8^ |
| EndB-m6 | No A domain | - | l-*allo*-Thr^8^ |
| ChersB-m6 | No A domain | - | l-*allo*-Thr^8^ |
| *A. orientalis* B-37-m6 | No A domain | - |  |
| *A. orientalis* DSM 40040-m6 | No A domain | - |  |
| *A. balhimycina*-m6 | No sequencing data | - |  |
| *Streptomyces* sp. TLI-053-m6 | No A domain | - |  |
| NRPS 2 m7 |  |  |  |
| RamoB-m7 | DAWTVAAV | l-Phe | l-Phe^9^ |
| EndB-m7 | DMEADGAV | l-hydrophillic | l-Cit^9^ |
| ChersB-m7 | DAWTVAAV | l-Phe | l-Phe^9^ |
| *A. orientalis* B-37-m7 | DAWTVAAV | l-Phe |  |
| *A. orientalis* DSM 40040-m7 | DAWTVAAV | l- Phe |  |
| *A. balhimycina*-m7 | No sequencing data | l- Phe |  |
| *Streptomyces* sp. TLI-053-m7 | DAWTVAAV | l- Phe |  |
| NRPS 3 m1 |  |  |  |
| RamoC-m1 | DMDTDGSV | d-/unknown | d-Orn^10^ |
| EndC-m1 | DAETDGSV | d-Orn, Lys, Arg | d-End^10^ |
| ChersC-m1 | DMETDGSV | d-Orn, Lys, Arg | d-Orn^10^ |
| *A. orientalis* B-37-m1 | DMETDGSV | d-Orn, Lys, Arg |  |
| *A. orientalis* DSM 40040-m1 | DMETDGSV | d-Orn, Lys, Arg |  |
| *A. balhimycina*-m1 | DMETDGSV | d-Orn, Lys, Arg |  |
| *Streptomyces* sp. TLI-053-m1 | DMETLVSV | d-Orn, Lys, Arg |  |
| NRPS 3 m2 |  |  |  |
| RamoC-m2 | DAFHLGLL | l-Hpg | l-Hpg^11^ |
| EndC-m2 | DAYHLGML | l-Hpg | l-Hpg^11^ |
| ChersC-m2 | DAYHLGLL | l-Hpg | l-Hpg^11^ |
| *A. orientalis* B-37-m2 | DAYHLGLL | l-Hpg |  |
| *A. orientalis* DSM 40040-m2 | DAYHLGLL | l-Hpg |  |
| *A. balhimycina*-m2 | DAYHLGML | l-Hpg |  |
| *Streptomyces* sp. TLI-053-m2 | DAYHLGLL | l-Hpg |  |
| NRPS 3 m3 |  |  |  |
| RamoC-m3 | DFWSVGMV | d-Thr | d-*allo*-Thr^12^ |
| EndC-m3 | DVWSVAMV | d-unknown | d-Ser^12^ |
| ChersC-m3 | DFWSVGMV | d-Thr | d-*allo*-Thr^12^ |
| *A. orientalis* B-37-m3 | DFWSVGMV | d-Thr |  |
| *A. orientalis* DSM 40040-m3 | DFWSVGMV | d-Thr |  |
| *A. balhimycina*-m3 | DFWSVGMV | d-Thr |  |
| *Streptomyces* sp. TLI-053-m3 | DFWNVGMV | d-Thr |  |
| NRPS 3 m4 |  |  |  |
| RamoC-m4 | DAYHLGLL | l-Hpg | l-Hpg^13^ |
| EndC-m4 | DAYHLGLL | l-Hpg | l-DiClHpg^13^ |
| ChersC-m4 | DALSLGTV | l-Phe, Trp, Phg, Tyr, Bht | l-Dpg^13^ |
| *A. orientalis* B-37-m4 | DAYHLGLL | l-Hpg |  |
| *A. orientalis* DSM 40040-m4 | DAYHLGLL | l-Hpg |  |
| *A. balhimycina*-m4 | DAFHLGLL | l-Hpg |  |
| *Streptomyces* sp. TLI-053-m4 | DALSLGTV | l-Gly, Ala, Val, Leu, Ile, Abu, Iva |  |
| NRPS 3 m5 |  |  |  |
| RamoC-m5 | DILQLGLV | Gly | Gly^14^ |
| EndC-m5 | DILQLGLV | Gly | Gly^14^ |
| ChersC-m5 | DILQLGLV | Gly | Gly^14^ |
| *A. orientalis* B-37-m5 | DILQVGLV | Gly |  |
| *A. orientalis* DSM 40040-m5 | DILQLGLV | Gly |  |
| *A. balhimycina*-m5 | DILQLGLV | Gly |  |
| *Streptomyces* sp. TLI-053-m5 | DILQLGLV | Gly |  |
| NRPS 3 m6 |  |  |  |
| RamoC-m6 | DAFFYGAT | l-Ile | l-Leu^15^ |
| EndC-m6 | DAETDGSV | l- Orn, Lys, Arg | l-End^15^ |
| ChersC-m6 | DAFWLGGT | l-Val | l-Val^15^ |
| *A. orientalis* B-37-m6 | DAMLVGAV | l-Val, Leu, Ile, Abu, Iva |  |
| *A. orientalis* DSM 40040-m6 | DAMLVGAL | l-Val, Leu, Ile, Abu, Iva |  |
| *A. balhimycina*-m6 | DAMLVGAV | l-Val, Leu, Ile, Abu, Iva |  |
| *Streptomyces* sp. TLI-053-m6 | DALWLGGT | l-Val |  |
| NRPS 3 m7 |  |  |  |
| RamoC-m7 | DVFSVAIL | d-Ala | d-Ala^16^ |
| EndC-m7 | DIFQLALV | d-Gly, Ala | d-Ala^16^ |
| ChersC-m7 | DVFSVAIV | d-Ala | d-Ala^16^ |
| *A. orientalis* B-37-m7 | DMET-GTV | d-hydrophillic |  |
| *A. orientalis* DSM 40040-m7 | DMETDGTV | d-hydrophillic |  |
| *A. balhimycina*-m7 | DAYHLGLL | d-Hpg |  |
| *Streptomyces* sp. TLI-053-m7 | DAYHLGLL | d-Hpg |  |
| NRPS 3 m8 |  |  |  |
| RamoC-m8 | DAYHLGLL | l-Hpg | l-ClHpg^17^ |
| EndC-m8 | DAYHLGLL | l-Hpg | l-Hpg^17^ |
| ChersC-m8 | DAYHLGML | l-Hpg | l-ClHpg^17^ |
| *A. orientalis* B-37-m8 | DAYHLGLL | l-Hpg |  |
| *A. orientalis* DSM 40040-m8 | DAYHLGLL | l-Hpg |  |
| *A. balhimycina*-m8 | DAYHLGLL | l-Hpg |  |
| *Streptomyces* sp. TLI-053-m8 | DALILGTV | l-Gly, Ala, Val, Leu, Ile, Abu, Iva |  |
| NRPS 4 |  |  |  |
| RamoD | DFWNIGMV | l-Thr | l-*allo*-Thr^8^ |
| EndD | DFWSVGMV | l-Thr | l-*allo*-Thr^8^ |
| ChersD | DFWNIGMV | l-Thr | l-*allo*-Thr^8^ |
| *A. orientalis* B-37 | DFWSIGMV | l-Thr |  |
| *A. orientalis* DSM 40040 | DFWSIGMV | l-Thr |  |
| *A. balhimycina* | DFWSVGMV | l-Thr |  |
| *Streptomyces* sp. TLI-053 | DFWSVGMV | l-Thr |  |

The eight adenylation domain specificity-conferring sequences were identified and predictions for the encoded amino acid are based on antiSMASH consensus and NRPSPredictor2. d- or l- stereochemistry is predicted based on the presence of ^L^C_L_ or E/C domains following the adenylation domain indicated.

**Table S4. NMR spectroscopic data of chersinamycin**

| **Residue** | **NH** | **α** | **β** | **other** |
| --- | --- | --- | --- | --- |
| **Asn1** | 7.91 | 4.29 | 2.05, 1.74 | - |
| **hyAsn2** | 8.26 | 5.27 | 5.55 |  |
| **Hpg3** | 9.58 | 5.98 | - | b/f 7.34; c/e 6.88 |
| **Orn4** | 9.05 | 4.10 | 1.22, 1.08 | γ 1.37, δ 2.68, 2.47 |
| **Thr5** | 7.43 | 4.17 | 3.89 | γ 0.94 |
| **Hpg6** | 8.80 | 6.63 | - | b/f 6.52; c/e 6.19 |
| **Hpg7** | 8.80 | 5.27 | - | b/f 6.52; c/e 6.30 |
| **Thr8** | 8.13 | 3.56 | 3.76 | γ 0.59 |
| **Phe9** | 7.47 | 4.01 | 2.05, 1.75 | b/f 6.80; c/e 7.09; d 7.04 |
| **Orn10** | 7.60 | 4.81 | 1.91, 1.83 | γ 1.54; δ 2.88, 2.82 |
| **Hpg11** | 9.10 | 6.80 | - | b/f 7.18; c/e 6.75 |
| **Thr12** | 8.93 |  | 3.79 | γ 0.80 |
| **Dpg13** | 8.57 | 5.79 | - | b/f 6.09; d 6.04 |
| **Gly14** | 7.76 | 3.60, 2.94 | - | - |
| **Val15** | 8.33 | 3.66 | 1.69 | γ 0.72 |
| **Ala16** | 9.26 | 4.16 | 1.23 | - |
| **Chp17** | 7.65 | 4.76 | - | b 6.20; e 6.67; f 6.35 |
| **lipid** | HC^α^ 1.97, HC^β^ 1.30, HC^γ^ 1.04, HC^δ^ 0.95, HC^ε^ 1.04, HC^ζ^ 0.95, HC^η^ 1.30, CH_3_ 0.65 | | | |

**Table S5. List of calculated and observed b- and y-ions from MS/MS of acyclic chersinamycin**

| **b ions** | **calculated** | | **observed** | |  | **y ions** | **calculated** | | **observed** | |
| --- | --- | --- | --- | --- | --- | --- | --- | --- | --- | --- |
|  | **M+1** | **M+2** | **M+1** | **M+2** |  |  | **M+1** | **M+2** | **M+1** | **M+2** |
| 1 | 155.144 |  | 155.144 |  |  | 1 | 202.027 |  |  |  |
| 2 | 269.187 |  | 269.187 |  |  | 2 | 273.064 |  | 273.064 |  |
| 3 | 399.224 |  | 399.121 |  |  | 3 | 372.132 |  | 372.129 |  |
| 4 | 548.272 |  | 548.275 |  |  | 4 | 429.154 |  | 429.154 |  |
| 5 | 662.351 |  | 662.359 |  |  | 5 | 594.196 |  | 594.194 |  |
| 6 | 763.400 |  | 763.394 |  |  | 6 | 695.244 |  | 695.242 |  |
| 7 | 912.447 |  | 912.445 |  |  | 7 | 1168.397 |  |  |  |
| 8 | 1061.494 | 531.251 |  |  |  | 8 | 1282.476 | 641.742 |  |  |
| 9 | 1162.542 | 581.774 | 1162.517 |  |  | 9 | 1429.545 | 715.276 |  |  |
| 10 | 1309.615 | 655.309 | 1310.609 |  |  | 10 | 1530.593 | 765.800 |  |  |
| 11 | 1423.690 | 712.384 | 1423.693 |  |  | 11 | 1679.640 | 840.322 |  |  |
| 12 | 1896.843 | 948.925 |  |  |  | 12 | 1828.688 | 914.848 |  |  |
| 13 | 1997.891 | 999.950 |  | 999.902 |  | 13 | 1929.736 | 965.371 | 1929.748 |  |
| 14 | 2162.933 | 1082.476 |  |  |  | 14 | 2083.815 | 1022.913 |  |  |
| 15 | 2219.955 | 1110.983 |  |  |  | 15 | 2192.863 | 1097.437 |  |  |
| 16 | 2319.023 | 1160.517 |  |  |  | 16 | 2322.900 | 1162.456 |  |  |
| 17 | 2390.060 | 1196.035 |  |  |  | 17 | 2436.943 | 1219.477 |  |  |
| 18 | 2573.069 | 1287.540 |  |  |  | 7a | 1006.344 |  | 1006.347 |  |
| 12a | 1734.790 | 867.899 |  |  |  | 8a | 1120.423 | 560.716 |  |  |
| 13a | 1835.837 | 918.422 |  |  |  | 9a | 1267.492 | 633.746 |  |  |
| 14a | 2000.887 | 1001.445 |  |  |  | 10a | 1368.539 | 684.774 |  |  |
| 15a | 2057.902 | 1029.956 |  |  |  | 11a | 1517.587 | 759.297 |  |  |
| 16a | 2156.967 | 1079.490 |  |  |  | 12a | 1666.635 | 833.821 |  |  |
| 17a | 2228.007 | 1115.009 |  |  |  | 13a | 1767.683 | 884.345 | 1767.670 |  |
| a | 2428.019 | 1215.015 |  | 1215.022 |  | 14a | 1881.762 | 941.385 |  |  |
| 12b | 1572.737 | 786.872 |  |  |  | 15a | 2030.810 | 1016.410 |  |  |
| 13b | 1673.785 | 837.396 | 1673.785 |  |  | 16a | 2160.848 | 1081.429 |  |  |
| 14b | 1838.827 | 919.917 |  |  |  | 17a | 2274.891 | 1138.451 |  |  |
| 15b | 1895.849 | 948.428 |  |  |  | 7b | 844.292 |  | 844.295 |  |
| 16b | 1994.917 | 998.464 |  |  |  | 8b | 958.371 | 479.689 | 958.372 |  |
| 17b | 2065.954 | 1033.982 |  | 1033.981 |  | 9b | 1105.439 | 553.233 | 1105.434 |  |
| b | 2265.967 | 1133.988 |  | 1134.026 |  | 10b | 1205.479 | 603.243 |  |  |
|  |  |  |  |  |  | 11b | 1355.535 | 678.271 | 1355.530 |  |
|  |  |  |  |  |  | 12b | 1504.582 | 752.795 | 1504.582 |  |
|  |  |  |  |  |  | 13b | 1605.630 | 803.319 | 1605.639 |  |
|  |  |  |  |  |  | 14b | 1719.709 | 860.358 |  |  |
|  |  |  |  |  |  | 15b | 1868.757 | 934.882 |  |  |
|  |  |  |  |  |  | 16b | 1998.795 | 1000.403 |  | 1000.405 |
|  |  |  |  |  |  | 17b | 2112.838 | 1057.424 |  |  |

a: fragment with loss of one sugar; b: fragment with loss of two sugars

**Table S6. Retention times for FDAA derivatives of amino acid standards and chersinamycin hydrolysate**

|  | **l-AA-FDAA** | **d-AA-FDAA** | **hydrolysate** |
| --- | --- | --- | --- |
| **Thr** | 11.75 | 15.17 |  |
| ***allo*-Thr** | 12.27 | 13.53 | 12.37, 13.42 |
| **FDAA** | 12.31 | - | 12.37 |
| **Gly** | 12.853 | - | 13.03 |
| **Ala** | 14.73 | 17.67 | 17.71 |
| **Hpg (mono)** | 18.01 | 20.56 | 18.19, 20.43 |
| **Val** | 20.39 | 24.17 | 20.43 |
| **Orn (di)** | 25.75 | 24.10 | 24.35 |
| **Phe** | 24.71 | 24.34 | 24.67 |
| **Hpg (di)** | 31.29 | 34.54 | 31.29, 34.59 |
| **ClHpg (di)** | 34.08 | - | 33.75 |
| **Asn** | 10.71 | 10.90 |  |
| **Dpg (mono)** | 16.21 | 17.14 |  |
| **Dpg (di)** | 29.71 | 31.47 |  |

Retention times are reported in minutes.

**Table S7.** **Deduced functions of proteins within the defined BGC of *M.* strain DSM 44151.**

| **Orf** | **Protein Product** | **Length** | **Protein Name** |
| --- | --- | --- | --- |
| 28 | WP_091305478.1 | 330 | hypothetical protein |
| 29 | WP_091321305.1 | 412 | PLP-dependent aminotransferase family protein |
| 30 | WP_091321307.1 | 260 | enoyl-CoA hydratase |
| 31 | WP_091321309.1 | 425 | enoyl-CoA hydratase/isomerase family |
| 32 | WP_091321311.1 | 205 | enoyl-CoA hydratase |
| 33 | WP_091305480.1 | 384 | type III polyketide synthase |
| 34 | WP_091305483.1 | 339 | 4-hydroxyphenylpyruvate dioxygenase |
| 35 | WP_091321312.1 | 388 | aminohydrolase family protein |
| 36 | WP_091321314.1 | 639 | ABC transporter ATP-binding protein |
| 37 | WP_091305485.1 | 266 | alpha/beta hydrolase |
| 38 | WP_091305488.1 | 529 | MBL fold metallo-hydrolase |
| 39 | WP_091305490.1 | 90 | acyl carrier protein |
| **Chers A** | **WP_091305493.1** | **2133** | **amino acid adenylation domain-containing protein** |
| **Chers B** | **WP_091305496.1** | **6998** | **amino acid adenylation domain-containing protein** |
| **Chers C** | **WP_091305499.1** | **8746** | **amino acid adenylation domain-containing protein** |
| 43 | WP_091305502.1 | 231 | thioesterase |
| 44 | WP_091305505.1 | 286 | NAD(P)-dependent oxidoreductase |
| **Chers D** | **WP_091321316.1** | **898** | **amino acid adenylation domain-containing protein** |
| 46 | WP_091305507.1 | 209 | class I SAM-dependent methyltransferase |
| 47 | WP_091305509.1 | 178 | hypothetical protein |
| 48 | WP_091305512.1 | 468 | DUF2029 domain-containing protein |
| 49 | WP_091305514.1 | 531 | FAD-dependent oxidoreductase |
| 50 | WP_091305517.1 | 218 | DNA-binding response regulator |
| 51 | WP_091321318.1 | 359 | two-component sensor histidine kinase |
| 52 | WP_091305519.1 | 184 | hypothetical protein |
| 53 | WP_091305522.1 | 301 | ABC transporter ATP-binding protein |
| 54 | WP_091305525.1 | 584 | hypothetical protein |
| 55 | WP_091321320.1 | 73 | MbtH family protein |
| 56 | WP_091305529.1 | 59 | hypothetical protein |
| 57 | WP_091305532.1 | 442 | cation/H(+) antiporter |
| 58 | WP_091321322.1 | 127 | chorismate mutase |
| 59 | WP_091321324.1 | 633 | hypothetical protein |
| 60 | WP_091305536.1 | 352 | alpha-hydroxy-acid oxidizing enzyme |

Numerical assignment of ORFs are derived from the original 81 analyzed translated sequences surround NRPS B.

**Table S8**. **Deduced functions of proteins within the defined BGC of *A. orientalis* strain B37.**

| Orf | Protein Product | Length | Protein Name |
| --- | --- | --- | --- |
| 29 | WP_052674858.1 | 332 | transcriptional regulator |
| 30 | WP_083255282.1 | 357 | streptomycin biosynthesis protein |
| 31 | WP_044850641.1 | 287 | 4-hydroxyphenylpyruvate dioxygenase |
| 32 | WP_052674849.1 | 789 | Aminotransferase |
| 33 | WP_044850640.1 | 778 | penicillin acylase family protein |
| 34 | WP_044850639.1 | 500 | FAD-dependent oxidoreductase |
| 35 | WP_065912850.1 | 341 | transmembrane transport protein |
| 36 | WP_044850637.1 | 308 | ABC transporter ATP-binding protein |
| 37 | WP_083254982.1 | 650 | ABC transporter ATP-binding protein |
| 38 | WP_044850636.1 | 275 | alpha/beta hydrolase |
| 39 | WP_044850635.1 | 90 | acyl carrier protein |
| **40** | **WP_052674848.1** | **2091** | **non-ribosomal peptide synthetase** |
| **41** | **WP_065912851.1** | **7005** | **non-ribosomal peptide synthetase** |
| **42** | **WP_065912852.1** | **8696** | **non-ribosomal peptide synthetase** |
| 43 | WP_044850632.1 | 236 | thioesterase |
| 44 | WP_044850631.1 | 274 | NAD(P)-dependent oxidoreductase |
| **45** | **WP_083254983.1** | **861** | **amino acid adenylation domain-containing protein** |
| 46 | WP_044850630.1 | 221 | DNA-binding response regulator |
| 47 | WP_083254984.1 | 421 | sensor histidine kinase |
| 48 | WP_044850753.1 | 169 | hypothetical protein |
| 49 | WP_083254985.1 | 373 | hypothetical protein |
| 50 | WP_044850629.1 | 554 | acyl-CoA dehydrogenase |
| 51 | WP_065912853.1 | 576 | acyl-CoA dehydrogenase |
| 52 | WP_083254986.1 | 618 | hypothetical protein |
| 53 | WP_037306096.1 | 74 | MbtH family protein |

Numerical assignment of ORFs are derived from the original 81 analyzed translated sequences surround NRPS B.

**Table S9. Deduced functions of proteins within the defined BGC of *A. orientalis* strain DSM 40040.**

| Orf | Protein product | Length | Protein name |
| --- | --- | --- | --- |
| 8 | WP_037306096.1 | 74 | MbtH family protein |
| 9 | WP_081736289.1 | 618 | hypothetical protein |
| 10 | WP_081736299.1 | 567 | acyl-CoA dehydrogenase |
| 11 | WP_051173836.1 | 554 | acyl-CoA dehydrogenase |
| 12 | WP_081736300.1 | 679 | hypothetical protein (mannosyltransferase) |
| 13 | WP_037306386.1 | 169 | hypothetical protein |
| 14 | WP_081736290.1 | 421 | sensor histidine kinase |
| 15 | WP_037306097.1 | 221 | DNA-binding response regulator |
| **16** | **WP_081736301.1** | **859** | **amino acid adenylation domain-containing protein** |
| 17 | WP_037306099.1 | 274 | NAD(P)-dependent oxidoreductase |
| 18 | WP_037306100.1 | 236 | Thioesterase |
| **19** | **WP_051173837.1** | **8720** | **non-ribosomal peptide synthetase** |
| **20** | **WP_051173838.1** | **7005** | **non-ribosomal peptide synthetase** |
| **21** | **WP_051173839.1** | **2091** | **non-ribosomal peptide synthetase** |
| 22 | WP_051173840.1 | 90 | polyketide synthase |
| 23 | WP_051173841.1 | 275 | alpha/beta hydrolase |
| 24 | WP_037306101.1 | 650 | ABC transporter ATP-binding protein |
| 25 | WP_051173842.1 | 308 | ABC transporter ATP-binding protein |
| 26 | WP_037306103.1 | 341 | Transporter |
| 27 | WP_037306105.1 | 500 | FAD-dependent oxidoreductase |
| 28 | WP_037306106.1 | 778 | penicillin acylase family protein |
| 29 | WP_037306109.1 | 795 | aminotransferase |
| 30 | WP_037306110.1 | 357 | 4-hydroxyphenylpyruvate dioxygenase |
| 31 | WP_081736302.1 | 287 | streptomycin biosynthesis protein |
| 32 | WP_037306397.1 | 332 | transcriptional regulator |

Numerical assignment of ORFs are derived from the original 60 analyzed translated sequences surround NRPS B.

**Table S10.** **Deduced functions of proteins within the defined BGC of *A. balhimycina* FH 1894 strain DSM 44591.**

| Orf | Protein product | Length | Protein name |
| --- | --- | --- | --- |
| 29 | WP_020647576.1 | 340 | hypothetical protein |
| 30 | WP_084642014.1 | 298 | streptomycin biosynthesis protein |
| 31 | WP_020647578.1 | 349 | 4-hydroxyphenylpyruvate dioxygenase |
| 32 | WP_020647579.1 | 805 | hypothetical protein |
| 33 | WP_051183855.1 | 779 | penicillin acylase family protein |
| 34 | WP_026469635.1 | 500 | FAD-dependent oxidoreductase |
| 35 | WP_026469636.1 | 341 | hypothetical protein |
| 36 | WP_051183856.1 | 311 | ABC transporter ATP-binding protein |
| 37 | WP_084642200.1 | 613 | ABC transporter ATP-binding protein |
| 38 | WP_020647585.1 | 280 | hypothetical protein |
| 39 | WP_020647586.1 | 90 | acyl carrier protein |
| 40 | **WP_084642015.1** | **2108** | **amino acid adenylation domain-containing protein** |
| 41 | - | - | - |
| 42 | **WP_020638000.1** | **8715** | **non-ribosomal peptide synthetase** |
| 43 | WP_026468001.1 | 236 | thioesterase |
| 44 | WP_020638002.1 | 274 | NAD(P)-dependent oxidoreductase |
| 45 | **WP_051183728.1** | **861** | **amino acid adenylation domain-containing protein** |
| 46 | WP_020638004.1 | 221 | DNA-binding response regulator |
| 47 | WP_020638005.1 | 420 | sensor histidine kinase |
| 48 | WP_020638006.1 | 170 | hypothetical protein |
| 49 | WP_020638007.1 | 566 | acyl-CoA dehydrogenase |
| 50 | WP_020638008.1 | 586 | acyl-CoA dehydrogenase |
| 51 | WP_084641135.1 | 620 | hypothetical protein |
| 52 | WP_020638010.1 | 74 | MbtH family protein |

Numerical assignment of ORFs are derived from the original 81 analyzed translated sequences surround NRPS B. While partial sequences for NRPS B were identified, no protein accession belonging to this region is available due to assembly gaps.

**Table S11.** **Deduced functions of proteins within the defined BGC of *Streptomyces* TLI-053.**

| Orf | Protein product | Length | Protein name |
| --- | --- | --- | --- |
| 35 | WP_107452518.1 | 141 | hypothetical protein |
| 36 | WP_093864800.1 | 302 | transcriptional regulator |
| 37 | WP_093859903.1 | 365 | prephenate dehydrogenase/arogenate dehydrogenase family protein |
| 38 | WP_107452520.1 | 375 | hydroxyneurosporene methyltransferase |
| 39 | WP_093864801.1 | 266 | amidinotransferase |
| **40** | **WP_093859905.1** | **8761** | **non-ribosomal peptide synthetase** |
| **41** | **WP_093859906.1** | **7121** | **amino acid adenylation domain-containing protein** |
| **42** | **WP_093859907.1** | **2139** | **amino acid adenylation domain-containing protein** |
| 43 | WP_093859908.1 | 90 | acyl carrier protein |
| 44 | WP_093859909.1 | 578 | acyl-CoA dehydrogenase |
| 45 | WP_093859910.1 | 581 | acyl-CoA dehydrogenase |
| 46 | WP_093859911.1 | 588 | hypothetical protein |
| 47 | WP_093859912.1 | 69 | MbtH family protein |
| 48 | WP_093859913.1 | 527 | MBL fold metallo-hydrolase |
| 49 | WP_093859914.1 | 268 | enoyl-CoA hydratase |
| 50 | WP_093859915.1 | 432 | enoyl-CoA hydratase/isomerase family protein |
| 51 | WP_093859916.1 | 219 | enoyl-CoA hydratase |
| 52 | WP_093859917.1 | 369 | type III polyketide synthase |
| 53 | WP_093859918.1 | 815 | aminotransferase |
| 54 | WP_093859919.1 | 337 | 4-hydroxyphenylpyruvate dioxygenase |
| 55 | WP_093859920.1 | 266 | alpha/beta hydrolase |
| 56 | WP_093859921.1 | 654 | ABC transporter ATP-binding protein |
| 57 | WP_093859922.1 | 330 | hypothetical protein |
| 58 | WP_093859923.1 | 300 | ABC transporter ATP-binding protein |
| 59 | WP_093859924.1 | 72 | hypothetical protein |
| 60 | WP_093859925.1 | 361 | hypothetical protein |
| 61 | WP_093859926.1 | 222 | DNA-binding response regulator |
| **62** | **WP_093859927.1** | **988** | **amino acid adenylation domain-containing protein** |
| 63 | WP_093859928.1 | 274 | NAD(P)-dependent oxidoreductase |
| 64 | WP_093859929.1 | 236 | thioesterase |

Numerical assignment of ORFs are derived from the original 81 analyzed translated sequences surround NRPS B.

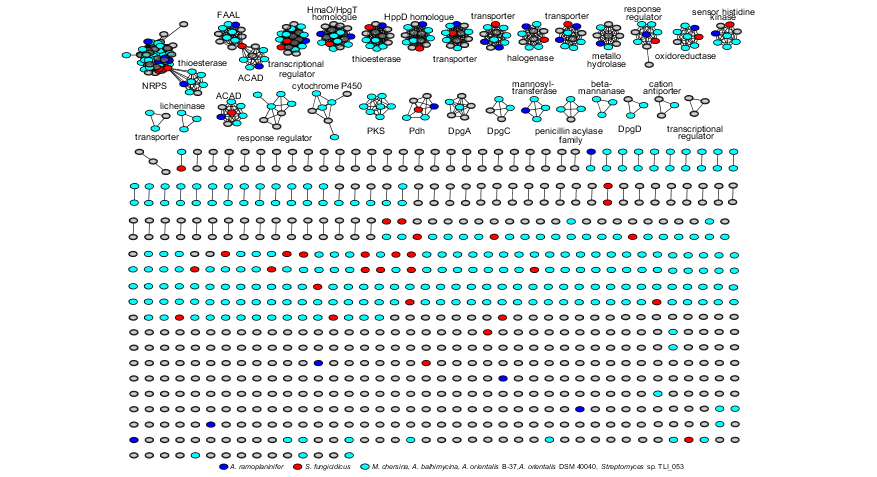

**Figure S1. Sequence similarity network of ORFs surrounding NRPS proteins in new bacterial strains.** The network is assembled for thirteen preliminary strains established through protein Blast analysis (listed in Table S1) with an E value limit of 10^-5^ and alignment score of 50. Proteins belonging to strains that were carried forward in further bioinformatic analyses are indicated in teal.

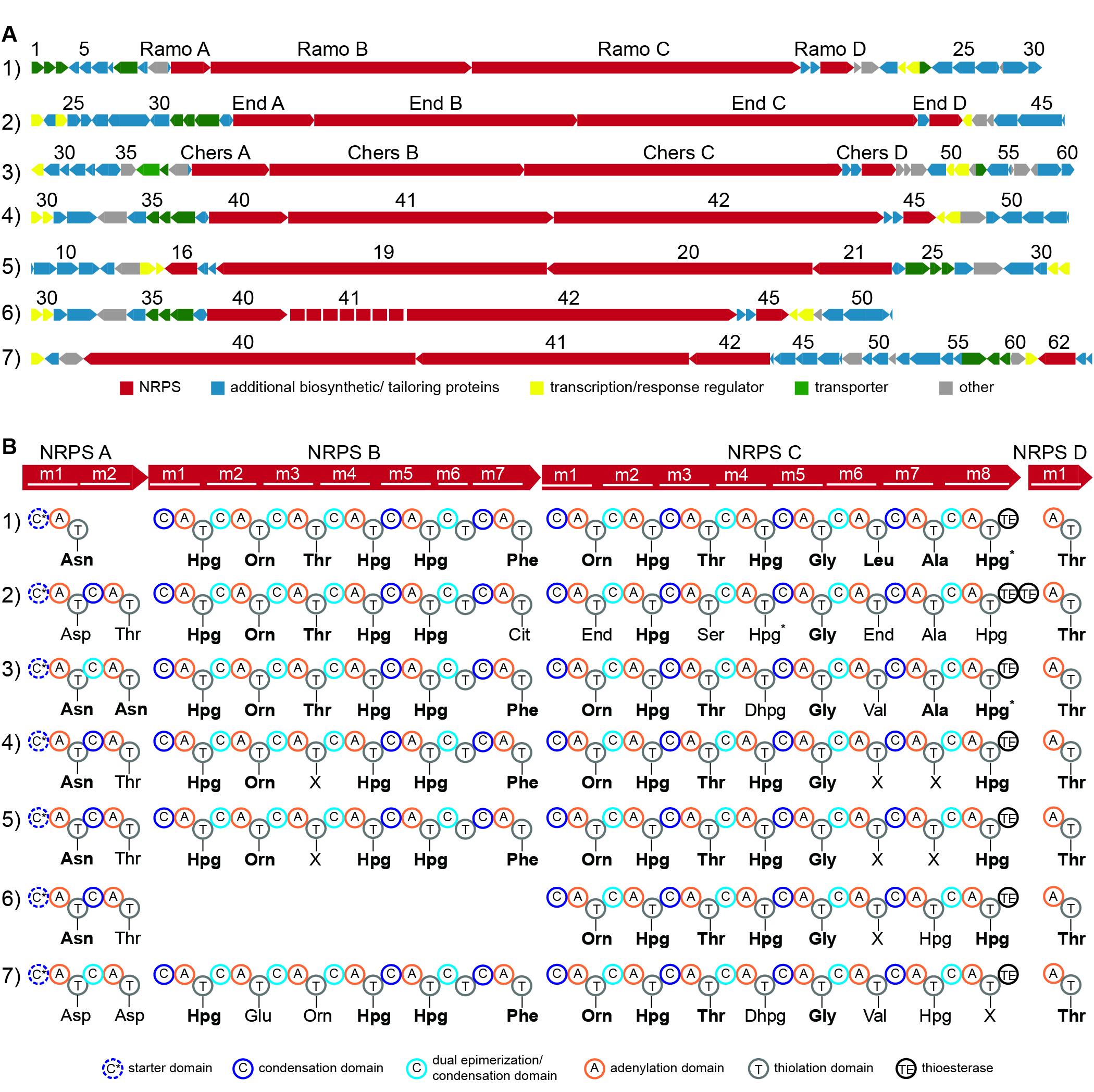

**Figure S2. (A) Open reading frame and (B) NRPS domain comparisons between ramoplanin family gene clusters.** (1) *A. ramoplanifer* strain ATCC 33076 (ramoplanin), (2) *S. fungicidicus* strain ATCC 21013 (enduracidin), (3) *M. chersina* strain DSM 44151 (chersinamycin), (4) *A. orientalis* strain B-37, (5) *A. orientalis* strain DSM 40040, (6) *A. balhimycina* strain FH189, and (6) *Streptomices* sp. TLI-053. Amino acids depicted for ramoplanin, enduracidin, and chersinamycin have been confirmed while those for the four remaining strains are based on predictions from conserved adenylation domain specificity sequences. Bolded residues highlight conserved residues relative to ramoplanin. Residues indicated with an “X” could not be predicted. An asterisk denotes a characterized chlorinated residue, though the adenylation domain confers specificity for Hpg.

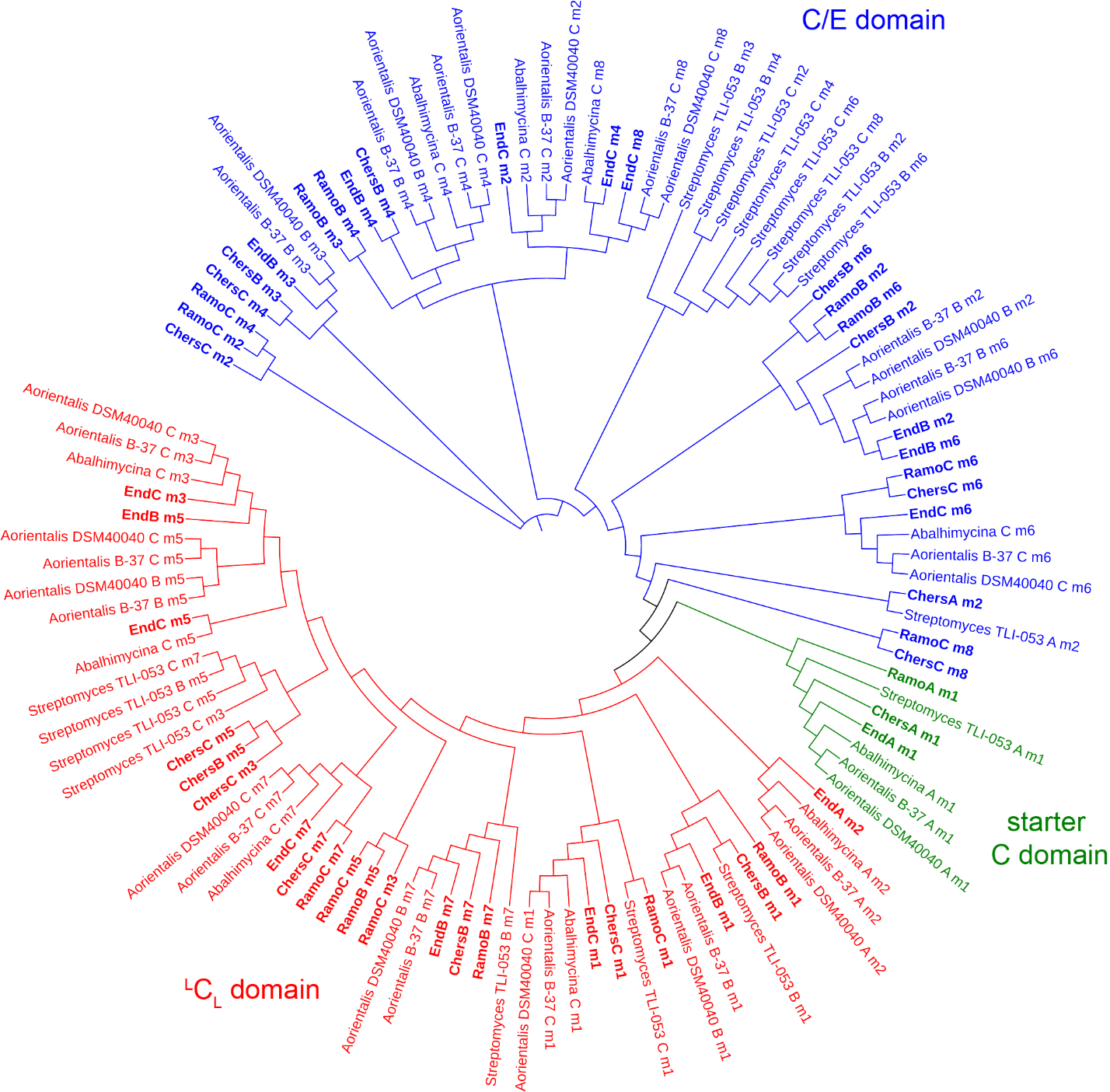

Figure S3. Phylogenetic relationship between NRPS condensation domains. Clusters are colored by C domain subtype: conventional ^L^C_L_ domains for l-amino acid incorporation, dual C/E domains for d-amino acid incorporation, and starter C domains for N-acyl lipid attachment. Domains in bold correspond to the C domains for characterized peptides ramoplanin, enduracidin, and chersinamycin.

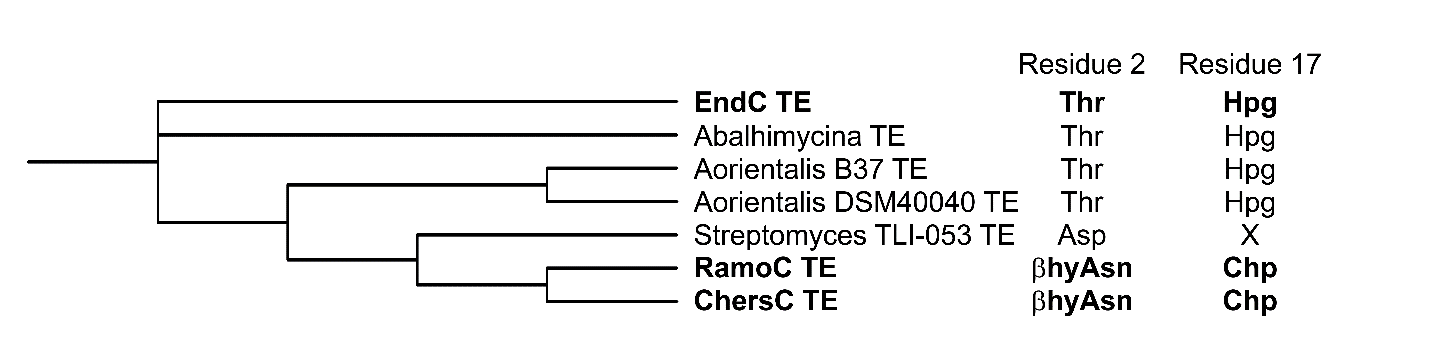
**Figure S4.** **Phylogenetic relationship between terminal NRPS C thioesterase domains**. Bolded letters indicate confirmed amino acids in enduracidin, ramoplanin, and chersinamycin.

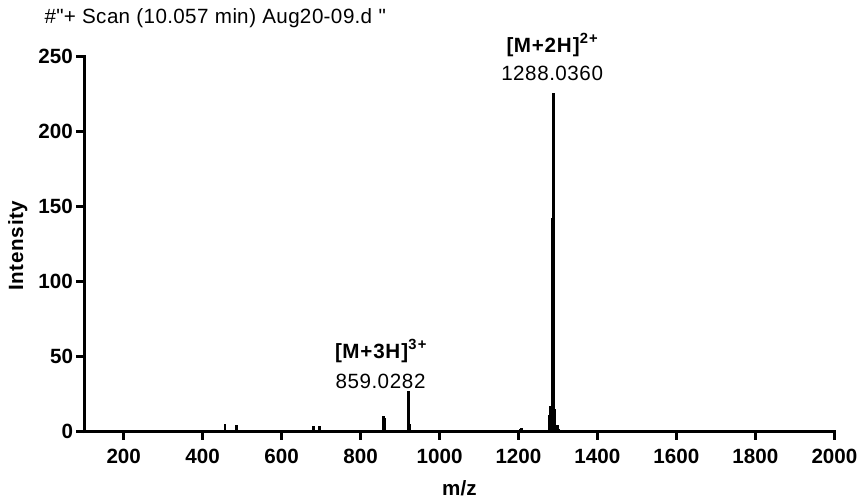

**Figure S5.** **HR-ESI-MS of chersinamycin.**

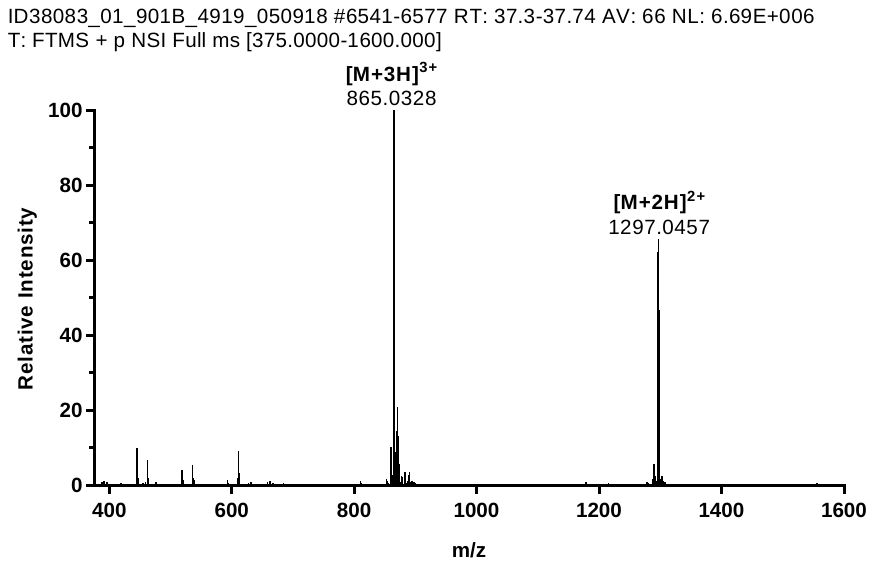

**Figure S6. HR-ESI-MS of acyclic chersinamycin.**

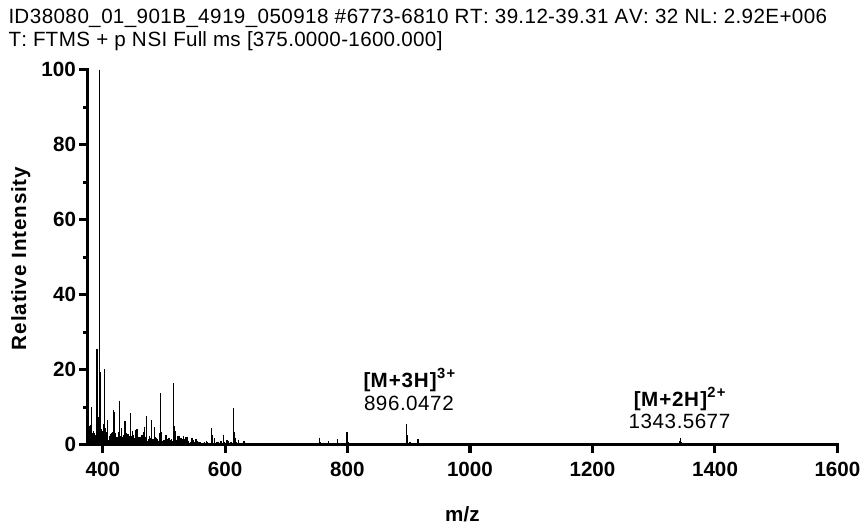

**Figure S7. ESI-MS spectrum of propionylated-ornithine-chersinamycin.**

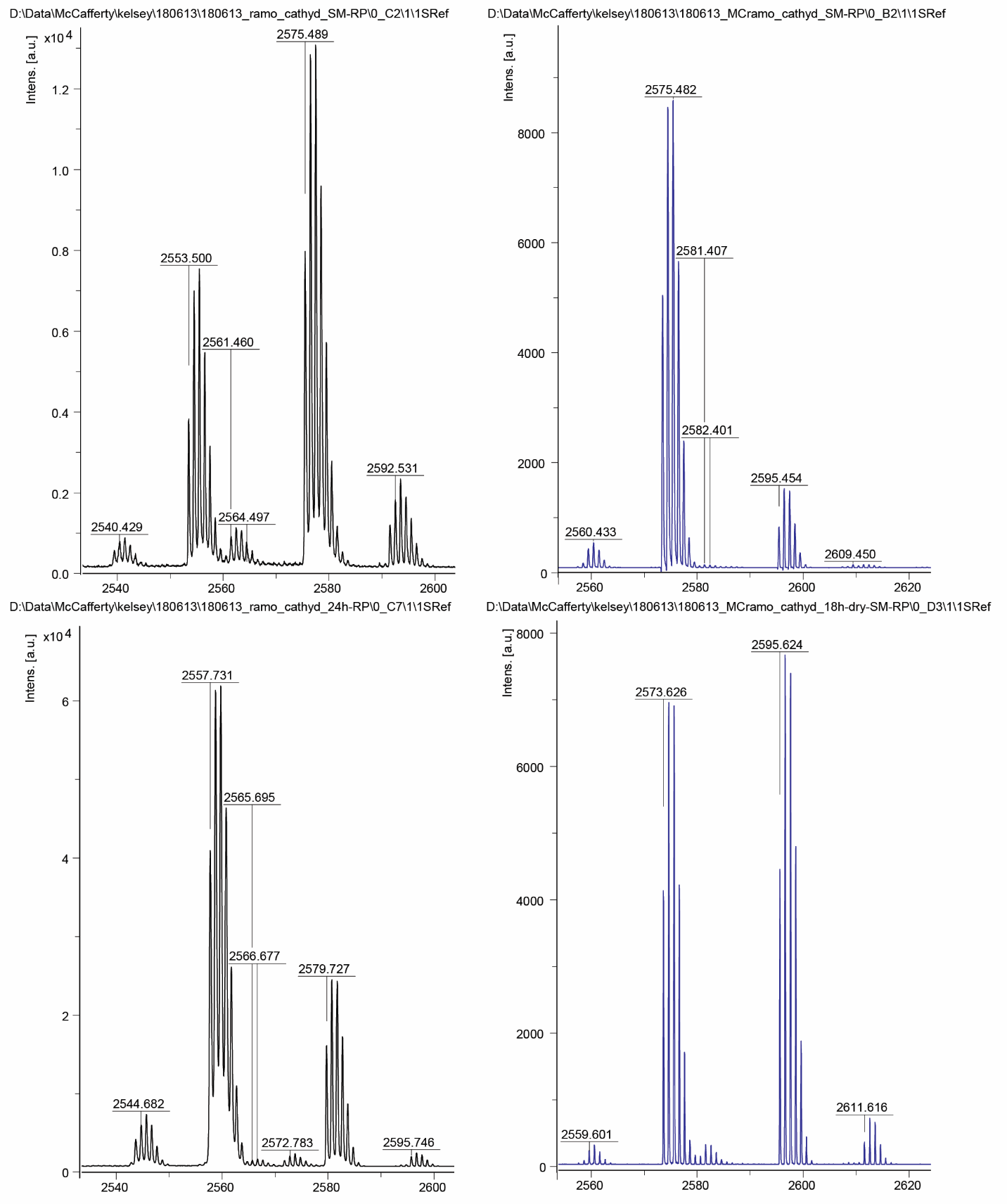

**Figure S8. MALDI-MS spectrum of hydrogenated ramoplanin (left) and chersinamycin (right).** The mass spectrum of hydrogenated ramoplanin (bottom) exhibits a clear 4 Da shift from starting material (top). The mass spectra for chersinamycin starting material (top) and hydrogenated product (bottom) are identical suggesting a saturated N-acyl lipid.

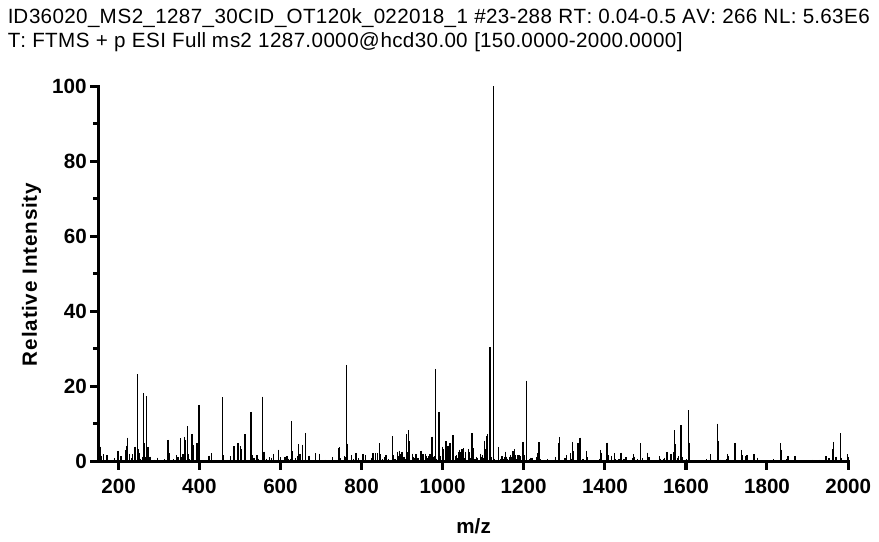

**Figure S9.** **ESI-MS/MS spectrum of chersinamycin.**

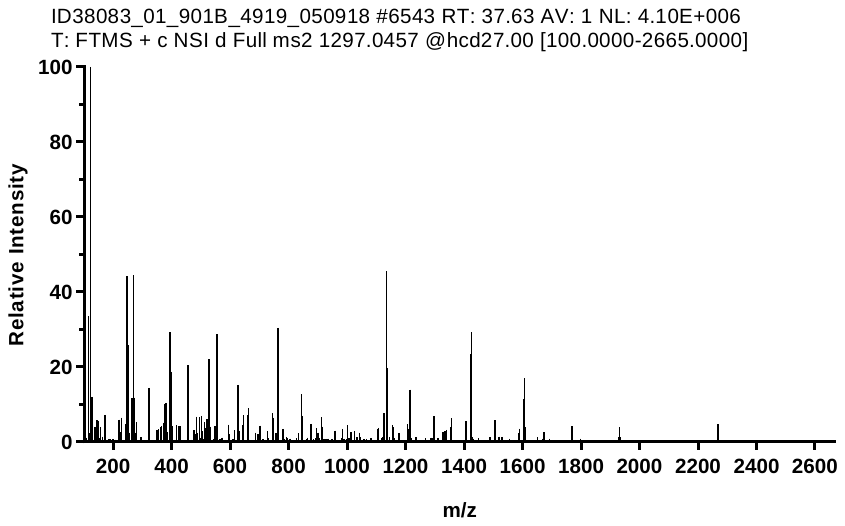

**Figure S10. ESI-MS/MS spectrum of acyclic chersinamycin**.

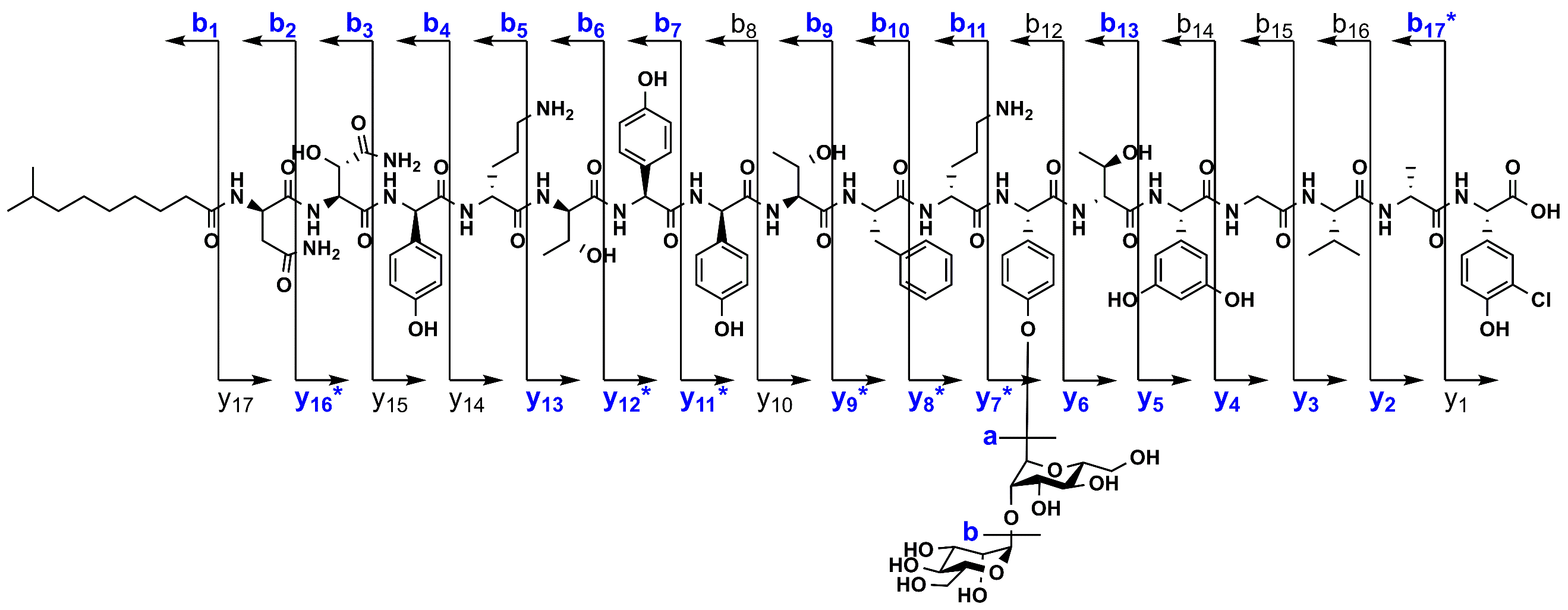

**Figure S11**. MS/MS fragmentation of acyclic chersinamycin (b- and y-ion series). The observed ions are shown in blue. An asterisk denotes fragments that were only observed with the loss of sugar units.

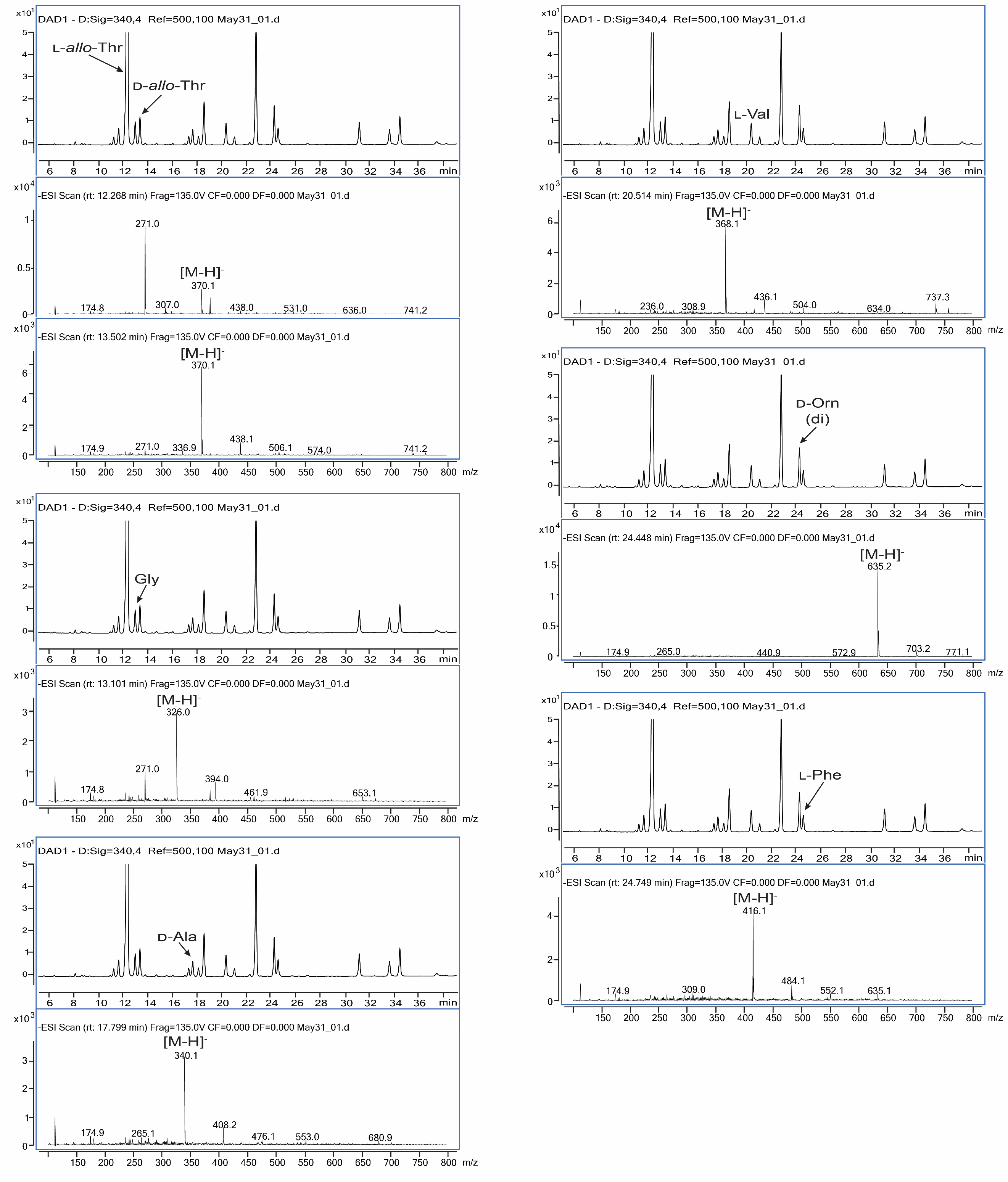

**Figure S12. Determination of absolute configuration of amino acids by advanced Marfey’s analysis.**

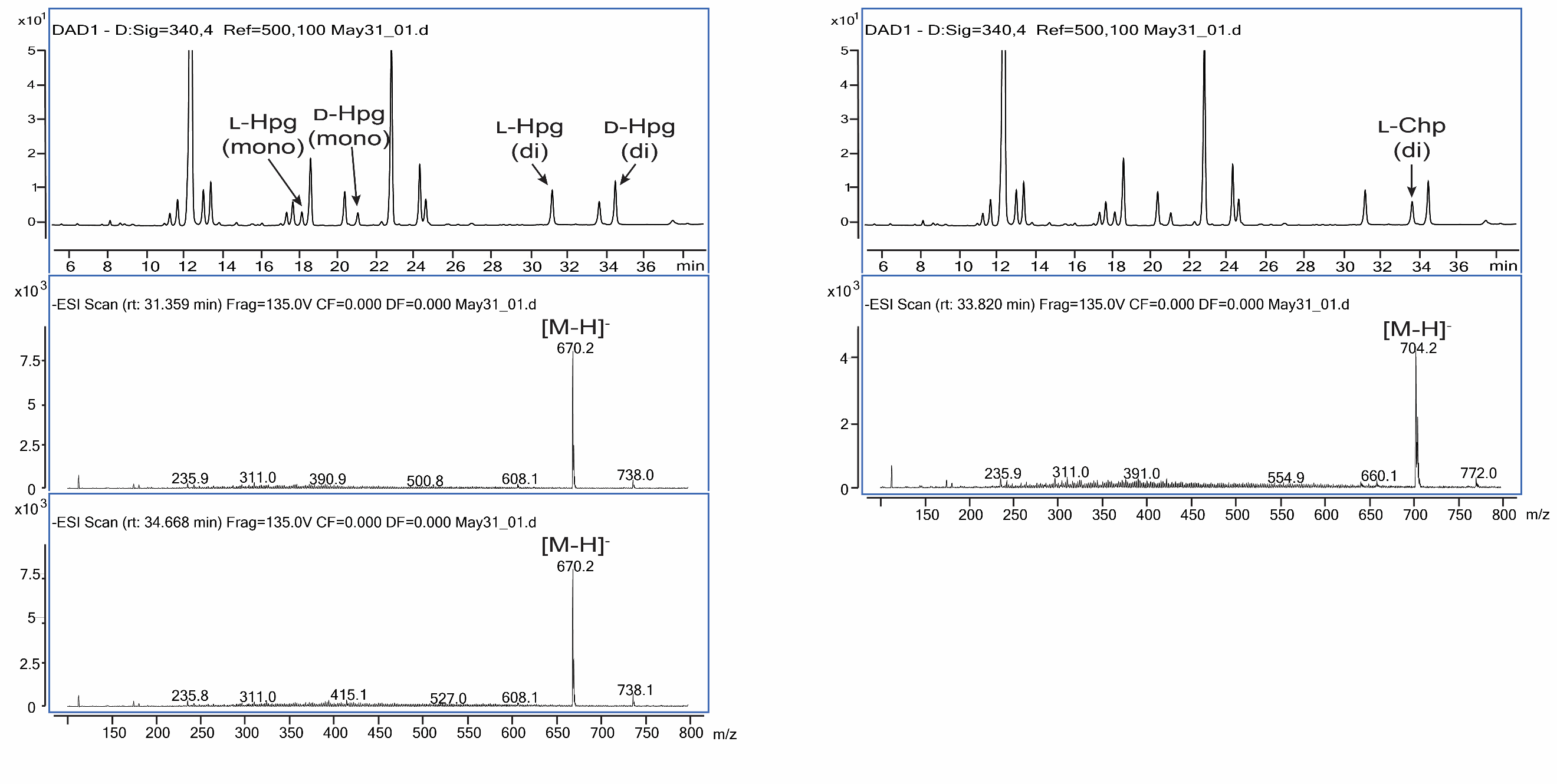

**Figure S12. Determination of absolute configuration of amino acids by advanced Marfey’s analysis (cont.).**

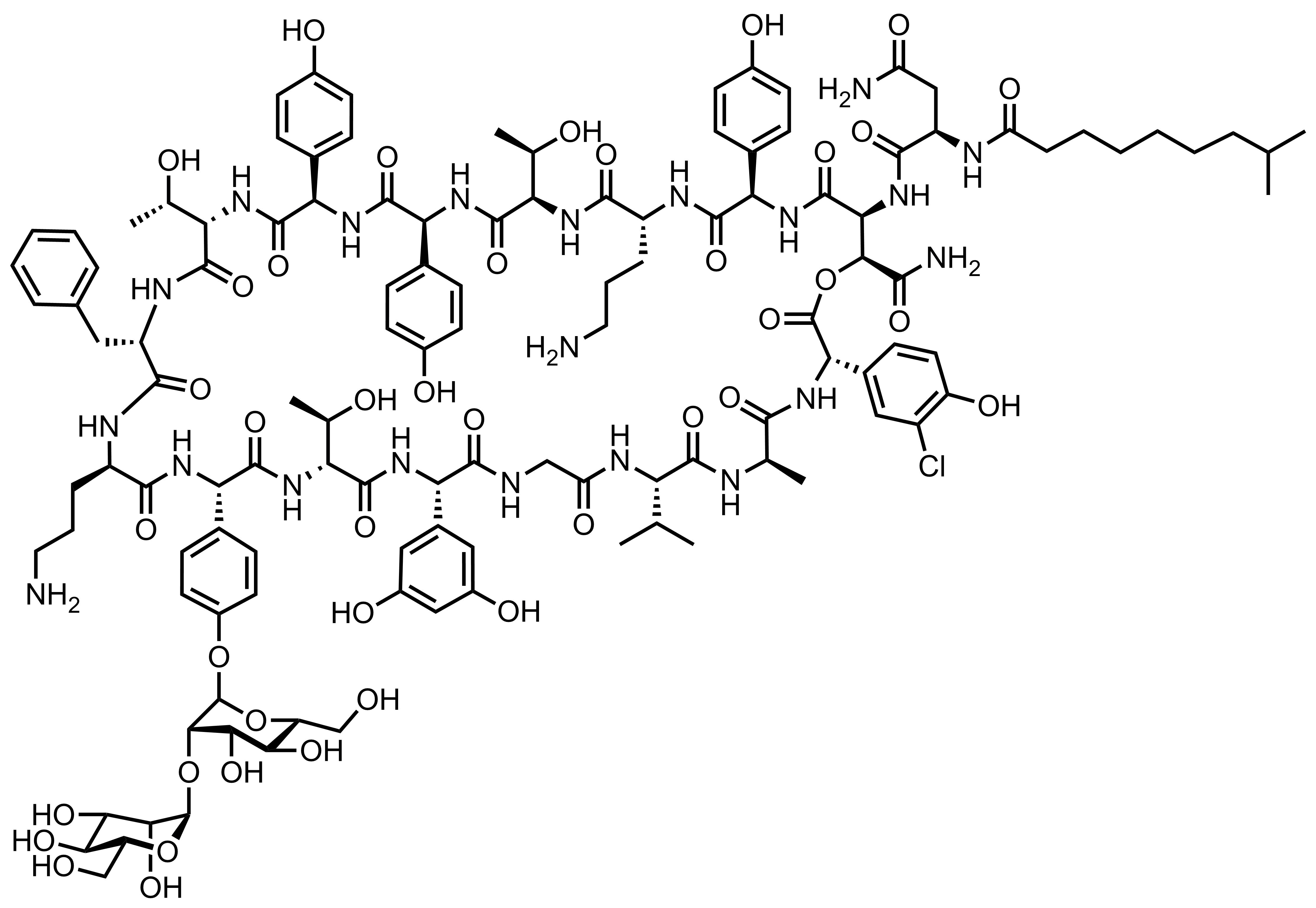

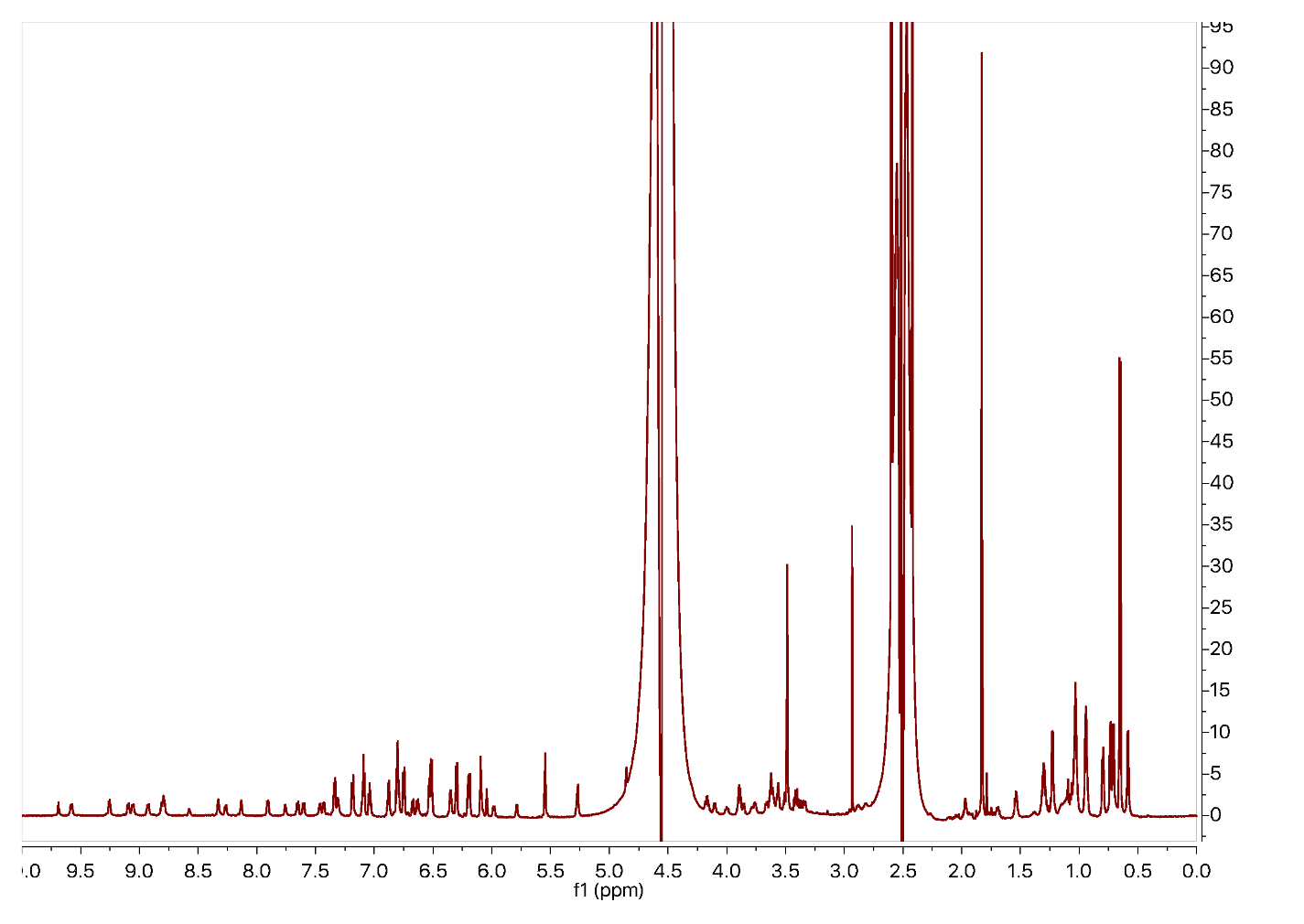

**Figure S13. ^1^H NMR (800 MHz, 4:1 H_2_O/DMSO-*d_6_*) spectrum of chersinamycin**.

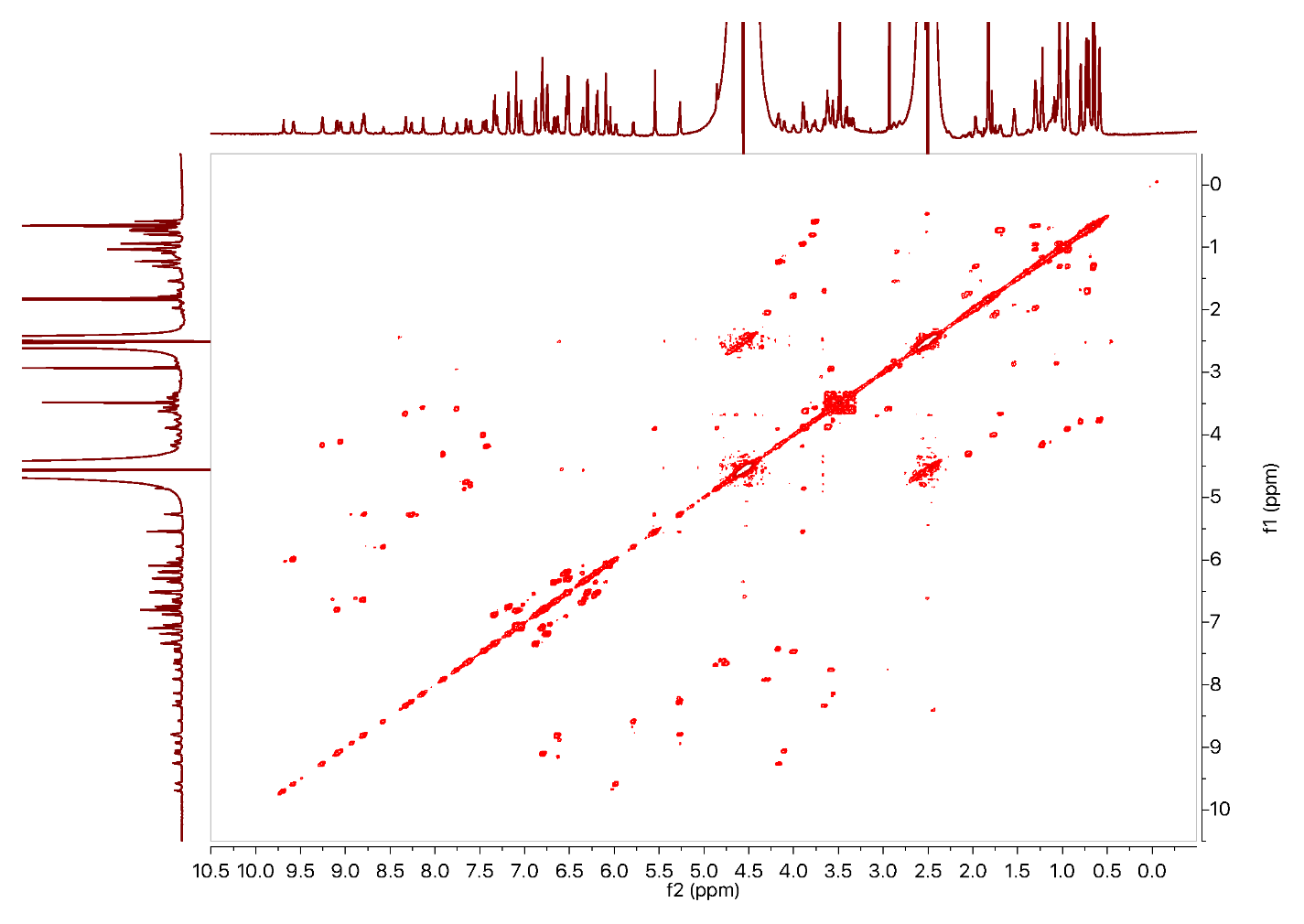

**Figure S14. ^1^H-^1^H COSY (800 MHz, 4:1 H_2_O/DMSO-*d_6_*) spectrum of chersinamycin.**

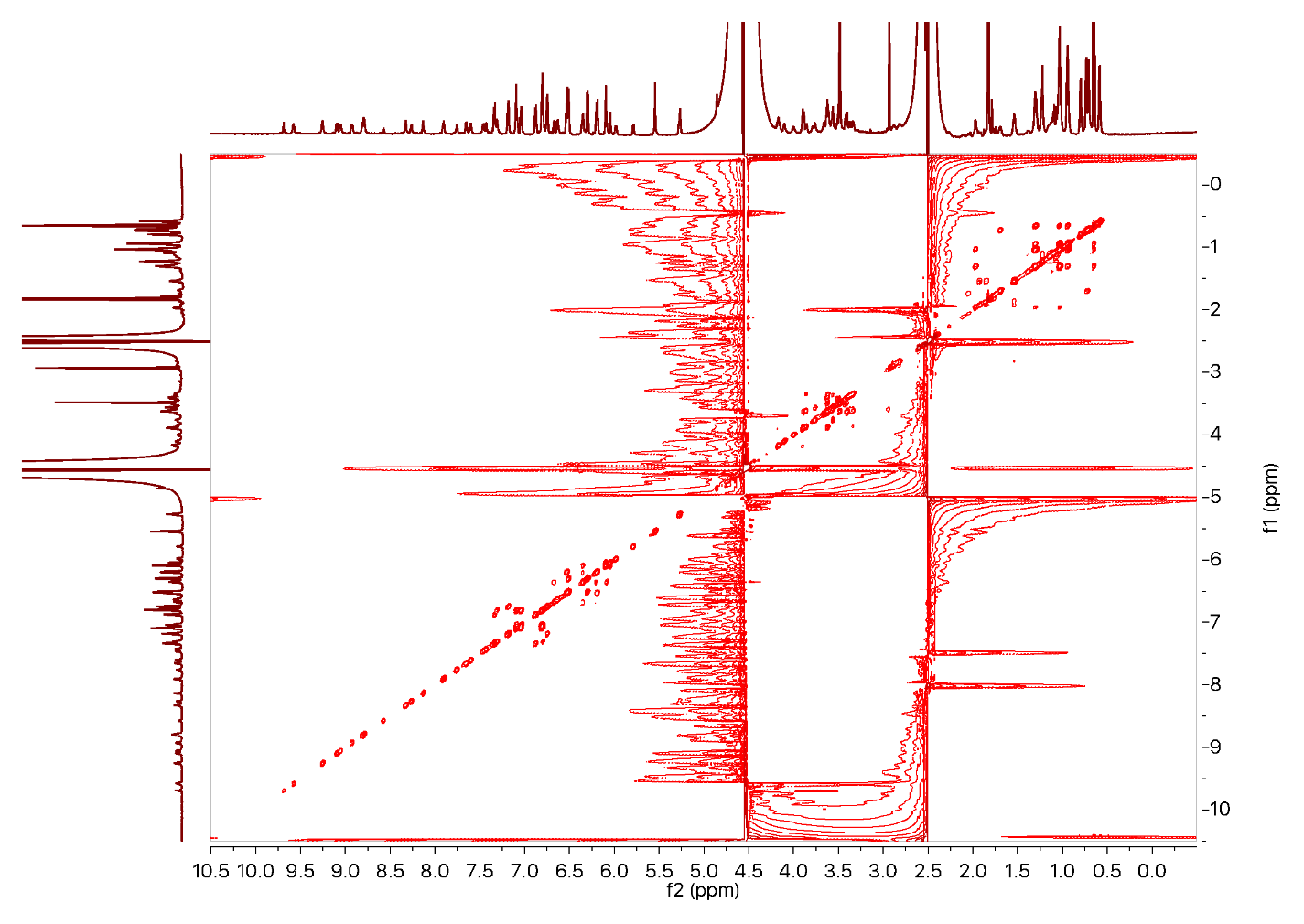

**Figure 15. ^1^H-^1^H TOCSY (800 MHz, 4:1 H_2_O/DMSO-*d_6_*) spectrum of chersinamycin.**

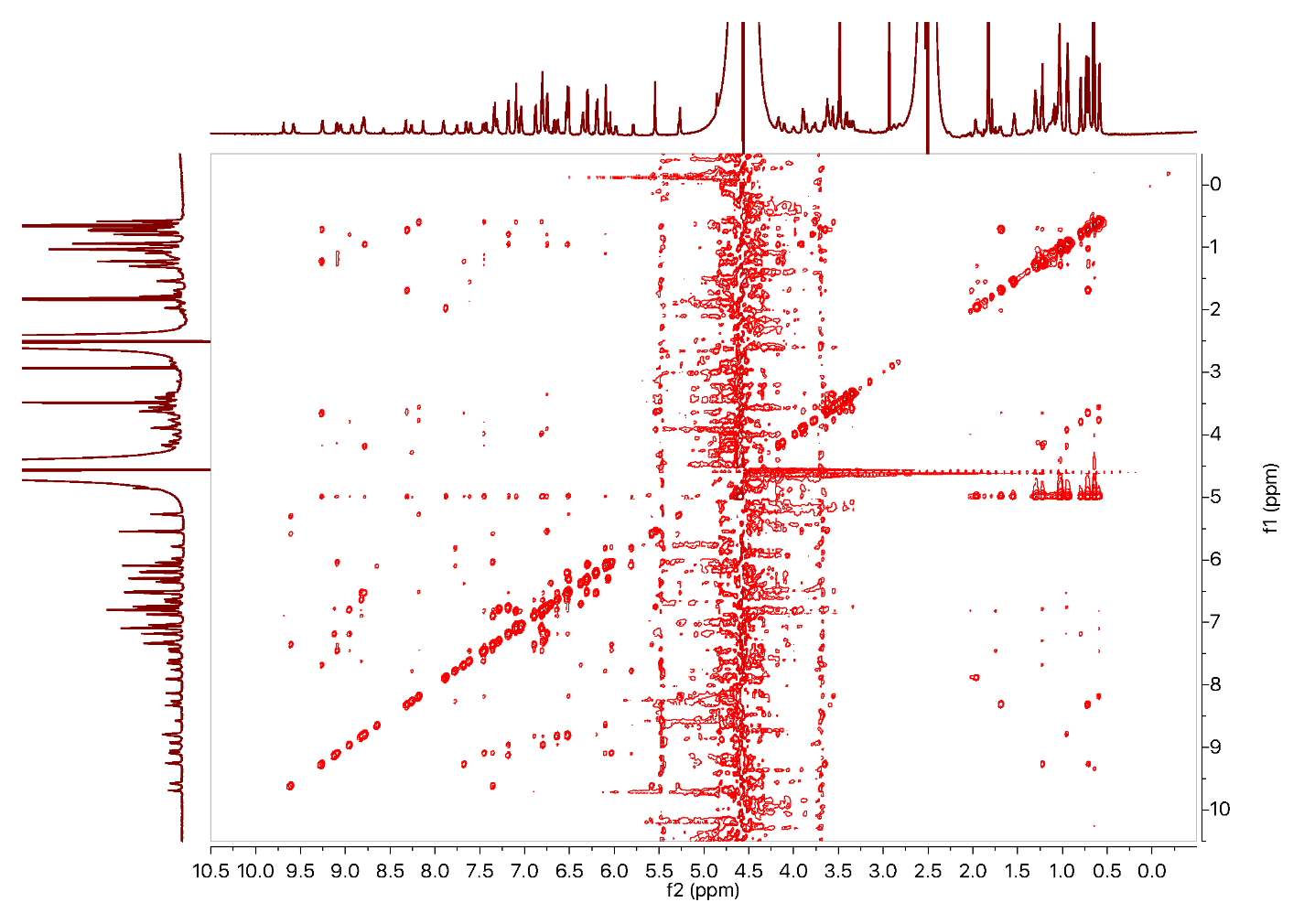

**Figure 16. ^1^H-^1^H NOESY (800 MHz, 4:1 H_2_O/DMSO-*d_6_*) spectrum of chersinamycin.**

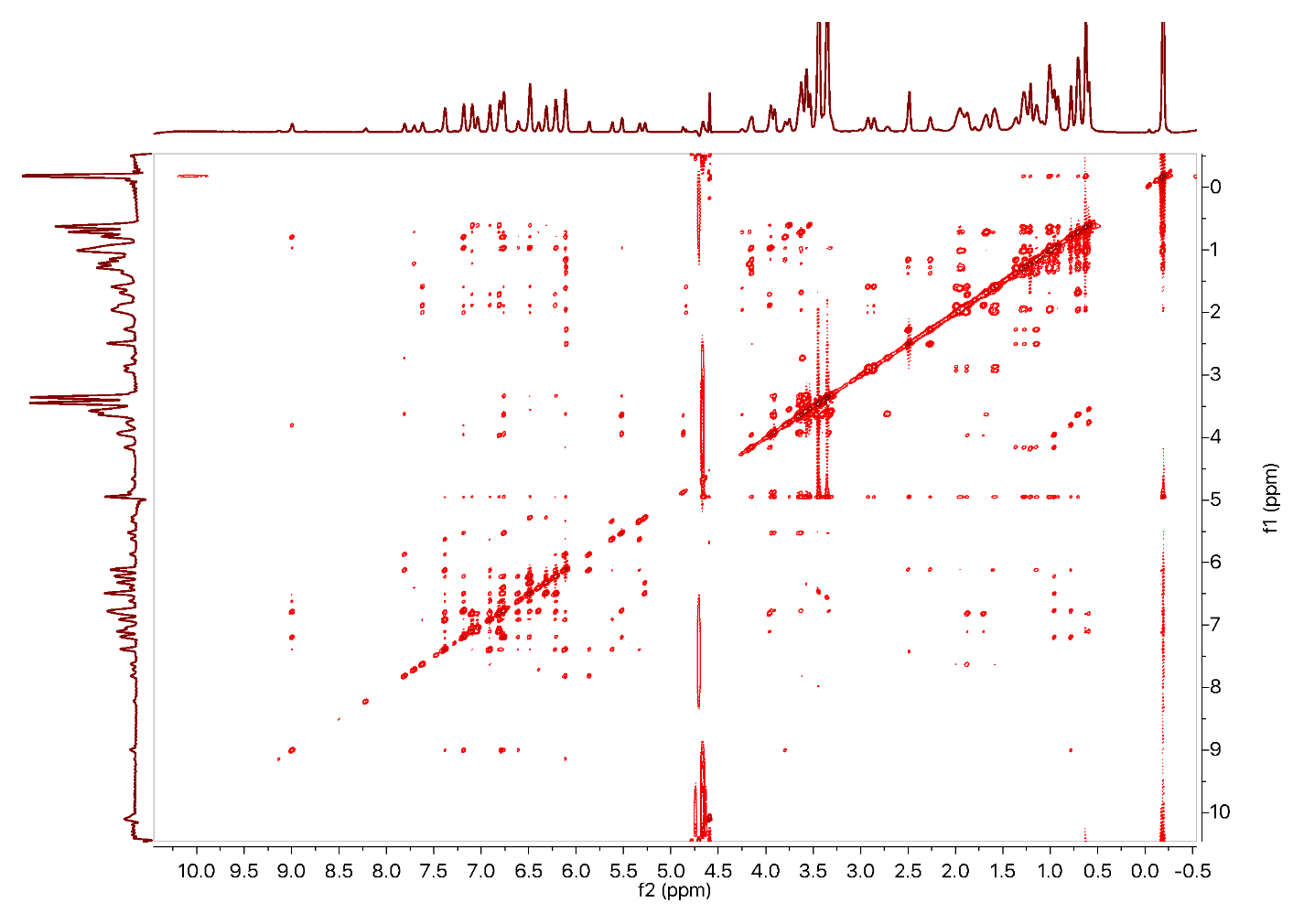

**Figure S17. ^1^H-^1^H NOESY (800 MHz, D_2_O/DMSO-*d_6_*) spectrum of chersinamycin.**

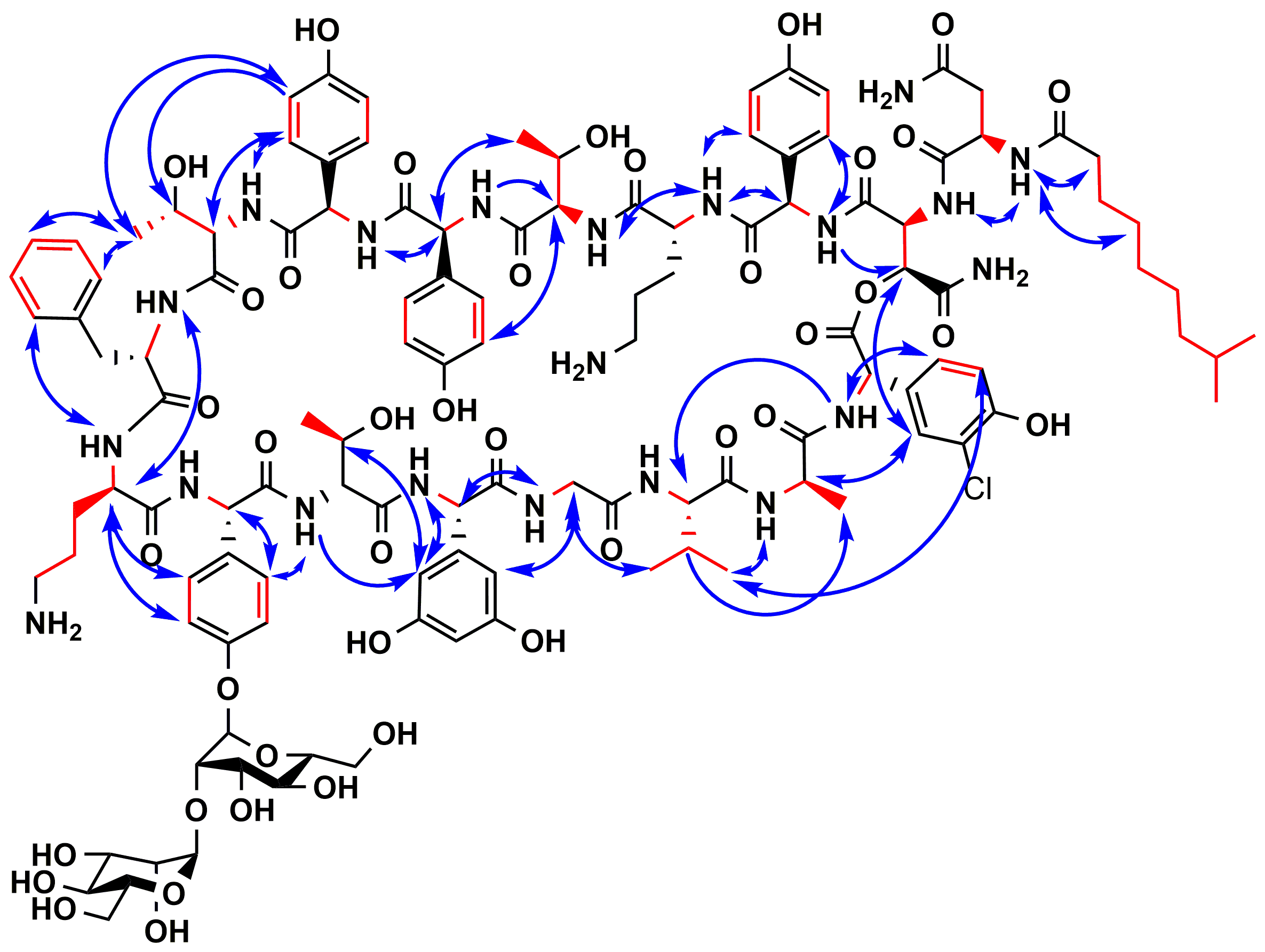

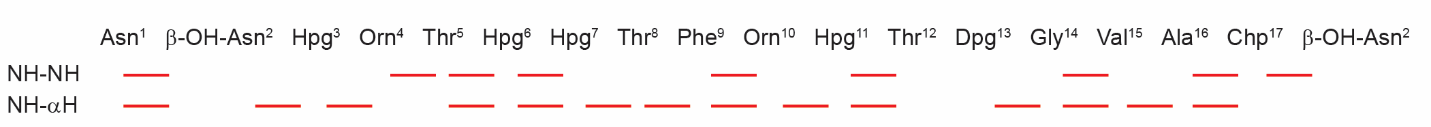

**Figure S18.** **Depiction of defining NMR correlations observed in chersinamycin**. COSY/TOCSY correlations are shown on the skeletal structure in red, and NOEs are depicted in blue. The inter-residue NOEs between adjacent amide protons (NH-NH) and adjacent amide and alpha protons (NH-αH) that were used to help determine connectivity are highlighted below the compound structure.
